# Divergent Molecular Pathways for Toxicity of Selected Mutant C9ORF72-derived Dipeptide Repeats

**DOI:** 10.1101/2023.09.28.558663

**Authors:** Sonia Okekenwa, Ming Ying Tsai, Patrick Dooley, Bin Wang, Priscila Comassio, Jorge E. Moreira, Nicola Kriefall, Sarah Y. Martin, Gerardo Morfini, Scott Brady, Yuyu Song

## Abstract

Expansion of a hexanucleotide repeat in a noncoding region of the C9ORF72 gene is responsible for a significant fraction of Amyotrophic Lateral Sclerosis (ALS) and Frontotemporal Dementia (FTD) cases, but mechanisms linking mutant gene products to neuronal toxicity remain debatable. Pathogenesis was proposed to involve the production of toxic RNA species and/or accumulation of toxic dipeptide repeats (DPRs) but distinguishing between these mechanisms has been challenging. In this study, we first use complementary model systems for analyzing pathogenesis in adult-onset neurodegenerative diseases to characterize the pathogenicity of DPRs produced by Repeat Associated Non-ATG translation of C9ORF72 in specific cellular compartments: isolated axoplasm and giant synapse from the squid. Results showed selective axonal and presynaptic toxicity of GP-DPRs, independent of associated RNA. These effects involved a MAPK signaling pathway that affects fast axonal transport and synaptic function, a pathogenic mechanism shared with other mutant proteins associated with familial ALS, like SOD1 and FUS. In primary cultured neurons, GP but not other DPRs promote the “dying-back” axonopathy seen in ALS. Interestingly, GR- and PR-DPRs, which had no effect on axonal transport or synaptic transmission, were found to disrupt the nuclear membrane, promoting “dying-forward” neuropathy. All C9-DPR-mediated toxic effects observed in these studies are independent of whether the corresponding mRNAs contained hexanucleotide repeats or alternative codons. Finally, C9ORF72 human tissues confirmed a close association between GP and active P38 in degenerating motor neurons as well as GR-associated nuclear damage in the cortex. Collectively, our studies establish compartment-specific toxic effects of C9-DPRs associated with degeneration, suggesting that two independent pathogenic mechanisms may contribute to disease heterogeneity and/or synergize on disease progression in C9ORF72 patients with ALS and/or FTD symptoms.

**Graphical Abstract:** **Activation of protein kinases and inhibition of axonal transport, synaptic transmission, and nuclear structure are toxic effects common to unrelated FALS-related gene products.** FALS-related mutant forms of SOD1 (mSOD1), FUS (mFUS), and C9-GP-DPRs (GP_(n)_) activate specific ASK1-MAPK pathway. Within axons, active ASK1-p38 pathway phosphorylates various substrates, including conventional kinesin, leading to the inhibition of fast axonal transport mediated by the translocation of this motor protein along microtubules. ASK1 can also inhibit synaptic transmission via JNK activation. Both pathways cause reductions in the availability of critical synaptic cargoes, synaptic dysfunction, and “dying-back” degeneration of neurons. On the other hand, C9- PR and GR-DPRs (PR_(n)_ and GR_(n)_) activate other pathways, leading to aberrant alterations in nuclear structure and function and “dying-forward” degeneration of neurons, consistent with reports of transcriptional changes and activation of apoptosis in ALS.

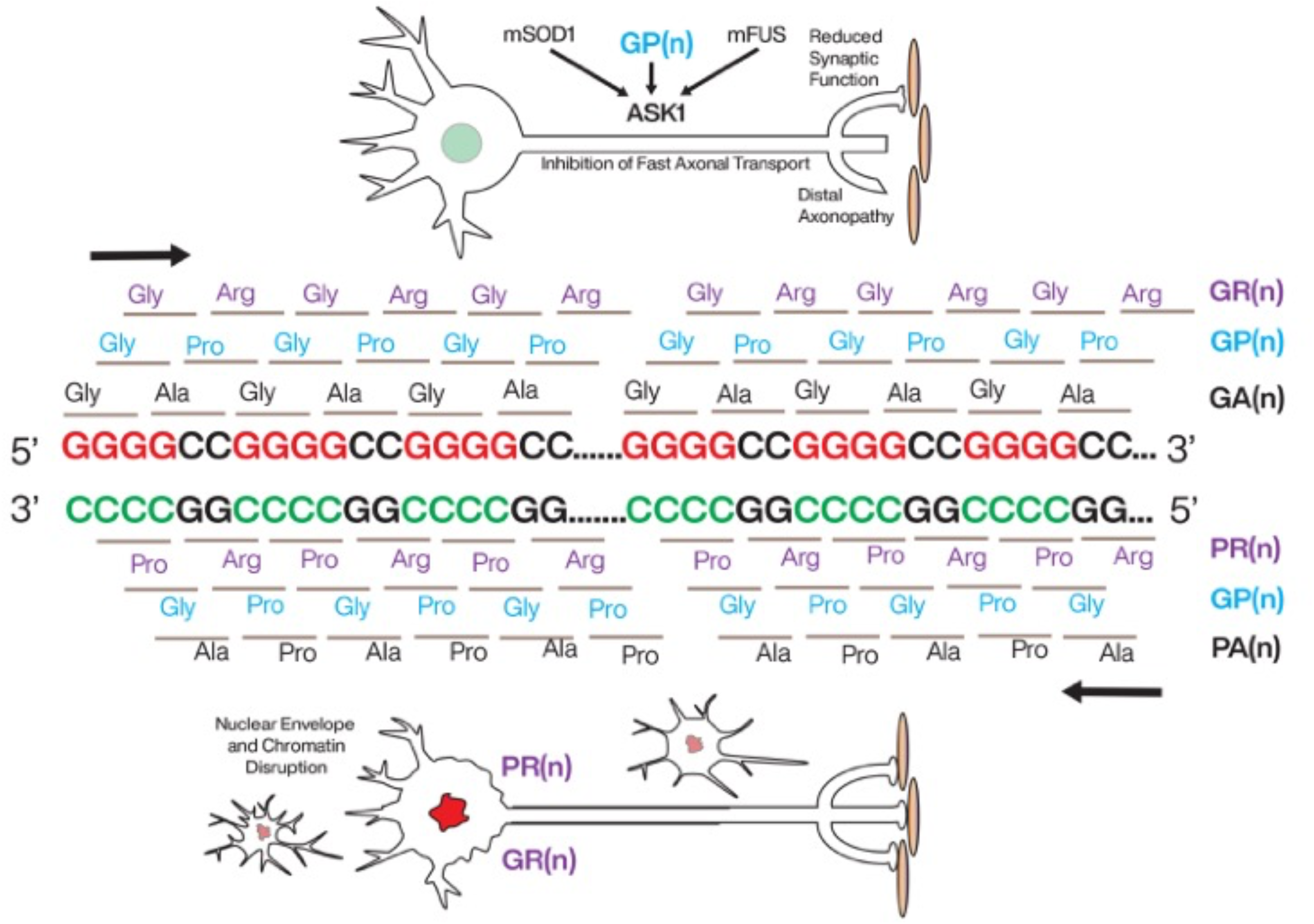

## INTRODUCTION

The surprising discovery that mutations in the C9ORF72 gene were involved in a significant number of both familial (∼40%) and sporadic (∼7%) Amyotrophic Lateral Sclerosis (ALS) cases, as well as ∼25% of Frontal Temporal Dementia (FTD) cases ^[1]^ presented a conundrum. The demonstration that the mutation involved expansion of a hexanucleotide repeat (GGGGCC) in a non-coding region of a gene of unknown function^[2, 3]^ added further complexity to the mystery. How do multiple mutations in a wide range of unrelated genes all render upper and lower motor neurons increasingly vulnerable in ALS^[4–6]^? Both loss of function and gain of function mechanisms have been proposed to explain pathogenesis in ALS, and C9ORF72 mutations were no exception^[7–9]^. Loss of C9ORF72 function associated with altered transcription or splicing has been suggested to alter immune function^[10]^ or autophagy pathways^[11–13]^. However, neither conditional nor ubiquitous deletions of the mouse C9ORF72 ortholog produces motor neuron degeneration^[7]^. Instead, total knockout of the gene produces a fatal autoimmune disease^[14]^. Many studies suggesting gain-of-function mechanisms have focused on RNA toxicity associated with the hexanucleotide repeats and based on the RNA foci observed in affected cells^[15, 16]^. Concurrently, other studies have implicated the accumulation of dipeptide repeat proteins (DPRs)^[17]^ produced by repeat-associated non-ATG (RAN) translation^[17–19]^. GP_n_ is produced by translation in both forward and reverse directions. GA_n_ and GR_n_ DPRs are generated by translation of expanded hexanucleotide sequences in the forward direction, whereas PR_n_ and AP_n_ DPRs are produced by translation in the reverse direction. These DPRs have been shown to accumulate in affected cells, including cell bodies, distal axons, and synaptic terminals^[20, 21]^. Still, others have suggested alterations on nucleocytoplasmic trafficking, which could be due to either mutant RNAs or DPRs^[22, 23]^. Although these mechanisms are not mutually exclusive, dissecting the relative contribution of these various mechanisms has been problematic, given the lack of model systems that can simultaneously distinguish these mechanisms. Moreover, the molecular basis for the increased vulnerability of motor neurons has remained elusive, making the identification of therapeutic targets challenging.

In this study, we first exploited two unique model systems for analyzing pathogenic mechanisms underlying adult-onset neurodegenerative diseases: isolated axoplasm from the squid giant axon^[24]^ and the squid giant synapse^[25–27]^. Using these systems, we characterized the pathogenicity of five DPRs associated with RAN translation of mutant C9ORF72 (GP, GA, GR, PA, and PR) in axons and synapses, major ALS-vulnerable neuronal compartments. Remarkably, GP, but not other DPRs elicited toxic effects on fast axonal transport, a cellular process critical for sustained axonal health known to be affected in ALS. As reported before with mutant SOD1^[28, 29]^, inhibition of an ASK1-P38 MAP kinase cascade mitigated this toxic effect. Given that mutant SOD1 also affects synaptic function^[30]^, the effect of DPRs in the presynaptic terminal of the squid giant synapse was also examined. In this model system, GP, but not GR, caused deficits in synaptic function that were also prevented by ASK1 inhibition.

Concurrently, we evaluated the toxicity of specific DPRs in mammalian models by transfection of rat primary cultured embryonic motor neurons. Consistent with results in squid models, expression of GP-DPR produced a striking axonopathy consistent with the “dying-back” pattern of degeneration observed in ALS-affected neurons^[31]^, which was reduced by inhibition of ASK1 and P38 MAP kinases. As seen in the squid models, neither GA nor PA DPRs affected fast axonal transport or cell viability.

In contrast with results obtained from isolated squid axons, two additional C9ORF72 DPRs (GR and PR) elicited cytotoxic effects on both fibroblasts and cultured motor neurons. Expression of GR and PR DPRs resulted in somal and nuclear pathology, rather than a distal axonopathy, and the effect was not blocked by inhibition of either ASK1 or P38 MAP kinase, suggesting a “dying-forward” pathology *via* a distinct molecular mechanism. The effects of GP, GR, and PR DPRs in cultured neurons were independent of whether the mRNA encoding these dipeptides contained hexanucleotide repeats or used alternative codons, which would not form atypical secondary structures of RNA, indicating that the observed toxicity was not due to RNA-based mechanisms.

Previous reports demonstrated that activation of P38 MAP kinase is a hallmark of both familial and sporadic forms of ALS^[32–36]^. Further, activation of ASK1 and P38 MAP kinase is a common feature of multiple gene products implicated in ALS pathology, including mutant and misfolded SOD1^[28, 29, 37, 38]^ and FUS^[39]^. Here, our results using ASK1 and P38 inhibitors suggest that, for C9ORF72 mutations, the toxicity of GP DPRs may be primarily responsible for motor neuron pathology *via* activation of an ASK1-MAPK pathway commonly shared by all other ALS-associated proteins examined to date. However, PR and GR DPRs may contribute to the development of neurodegeneration *via* an unknown pathway specific for patients with the expansion of the hexanucleotide repeat in C9ORF72. Finally, immunostaining of human tissues obtained from C9-ORF72-ALS patients revealed a close association of GP DPRs with active p38 kinases in affected motor neurons, while aggregated GR was associated with abnormal nuclear envelope in cortical neurons affected in C9-ORF72 FTD. Identification of two divergent pathogenic pathways elicited by specific C9-derived DRPs may help explain the heterogeneous diseases elicited by c9orf72 mutations.

## RESULTS

### GP dipeptide repeat protein inhibited axonal transport via the ASK1-P38 pathway

ALS proceeds as a dying back axonopathy with loss of distal axons and synapses as early pathological events, suggesting that the axonal compartment is a critical pathogenic target in ALS-affected neurons^[31]^. Previous studies documented inhibition of fast axonal transport by ALS-linked forms of FUS and SOD1 *via* activation of ASK1-P38 MAP kinase and phosphorylation of kinesin heavy chains^[28, 29, 37, 39]^, raising the question of whether axonal P38 MAP kinase activation is a general feature of additional familial ALS (fALS)-linked mutant gene products. C9ORF72 is the gene most frequently mutated in ALS^[1, 5]^, but it is unusual in being a hexanucleotide repeat expansion that can produce both multiple distinct dipeptide repeat proteins (DPRs) through RAN translation and abnormal RNA foci.

RAN translation of the hexanucleotide repeat in mutant C9ORF72 resulted in the accumulation of five different DPRs of various lengths (GP, GR, PR, GA, PA) in patient brains^[18, 40, 41]^. To evaluate the toxic effects of individual DPRs in the axonal compartment, we performed vesicle motility assays following perfusion of synthetic DPRs (16-mer peptides encompassing 8 dipeptide repeats) at the same concentration into isolated squid axoplasm (**Fig. 1**). This *ex vivo* model lacks nuclear transcription, protein synthesis machinery, and glia, allowing testing of axonal transport-specific mechanisms. Since the DPRs were added as peptides, there was no exogenous RNA that could generate RNA toxicity. The absence of a plasma membrane in the isolated squid axon preparation allows the introduction of neuropathogenic proteins and pharmacological agents at defined concentrations, and quantitative analysis of fast axonal transport rates of membrane-bounded organelles (MBOs) moving in both anterograde (kinesin 1-dependent) and retrograde (cytoplasmic dynein-dependent) directions^[24]^.

**Figure 1.**
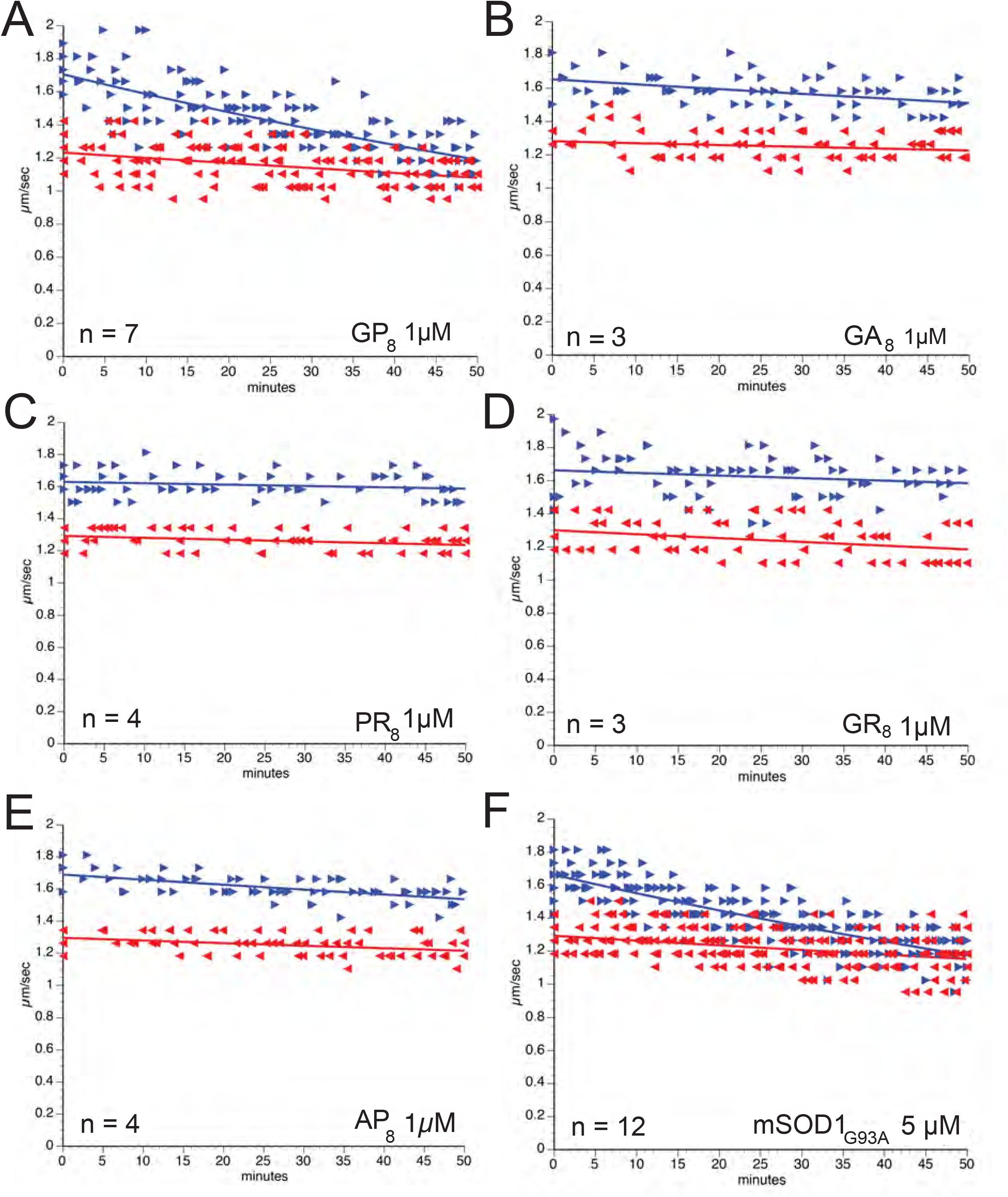
GP(n_8_) DPRs selectively inhibit anterograde axonal transport. Plots depict results from vesicle motility assays. Individual velocity (µm/sec) measurements (arrowheads) are plotted as a function of time (minutes) for a whole duration of 50min. Blue arrowheads and lines represent anterograde, kinesin-dependent fast axonal transport rates. Red reverse arrows and lines represent retrograde, cytoplasmic dynein-dependent axonal transport rates. Perfusion of GP(n_8_) peptide specifically inhibited anterograde, but not retrograde transport (**A**). In contrast, perfusions of GA(n_8_) (**B**), PR(n_8_) **(C)**, GR(n_8_) (**D**), or AP(n_8_) **(E)** peptides did not affect axonal transport in either direction. The toxic effect of GP(n_8_) was similar to those elicited by mutant SOD1-G93A (**F**), SOD1-H46R, and SOD1-G85R^[28, 29]^.

Among the five DPRs, only GP_8_ (Fig 1A) was found to affect vesicle motility in axoplasm by preferentially inhibiting anterograde transport, an effect similar to that observed for mutant SOD1(G93A and G85R)^[28, 29, 37]^ (Fig 1F) and FUS^[39]^. The effects of GA_8_, GR_8_, PR_8_, and AP_8_ (Fig 1B-E) were not different from buffer control, with no change in axonal transport rates observed for the time course of these experiments. Quantitative analysis of these data indicated that only GP_8_ and mutant SOD1(G93A) significantly inhibited the anterograde transport of vesicles (Fig 1).

Selective inhibition of anterograde axonal transport by GP_8_, an effect similar to that elicited by pathogenic forms of SOD1^[28, 29, 37]^ suggested that GP_8_ and ALS-linked forms of SOD1 may also activate a similar pathway. Pharmacological and biochemical experiments indicated that mutant SOD1 activates a MAP kinase signaling cascade culminating in phosphorylation of kinesin heavy chain by P38α MAPK^[28, 29]^. This cascade involved the activation of the MAP kinase kinase kinase (MAP3K) ASK1^[29, 39]^, which in turn activated a MAP kinase kinase (MAP2K), leading to p38 MAPK activation^[28]^. To determine whether toxic effects of GP_8_ on axonal transport were also mediated by p38α, we co-perfused axoplasms with GP_8_ and MW069, a selective inhibitor of P38α MAPK^[42]^. MW069 prevented inhibition of anterograde axonal transport by GP (Fig. 2A), as it did for mutant SOD1 (Fig. 2D), suggesting a critical role of P38α MAPK in axon-specific toxicity of GP. Co-perfusion of GP_8_ with a DVD peptide, which prevents a subset of MAP3Ks from docking to and activating downstream MAP2Ks^[43]^, also prevented the toxic effects of GP_8_ on anterograde axonal transport (Fig 2B). Similarly, co-perfusion with NQDI-1, an inhibitor of the MAP3K ASK1^[44]^, prevented inhibition of anterograde transport by GP_8_ (Fig 2C), as it did for mutant SOD1^[29]^ and FUS^[39]^. These studies suggest that pathological forms of SOD1, FUS, and C9ORF72-GP disrupt fast axonal transport through a common pathway: activation of the ASK1-P38 MAP kinase cascade.

**Figure 2.**
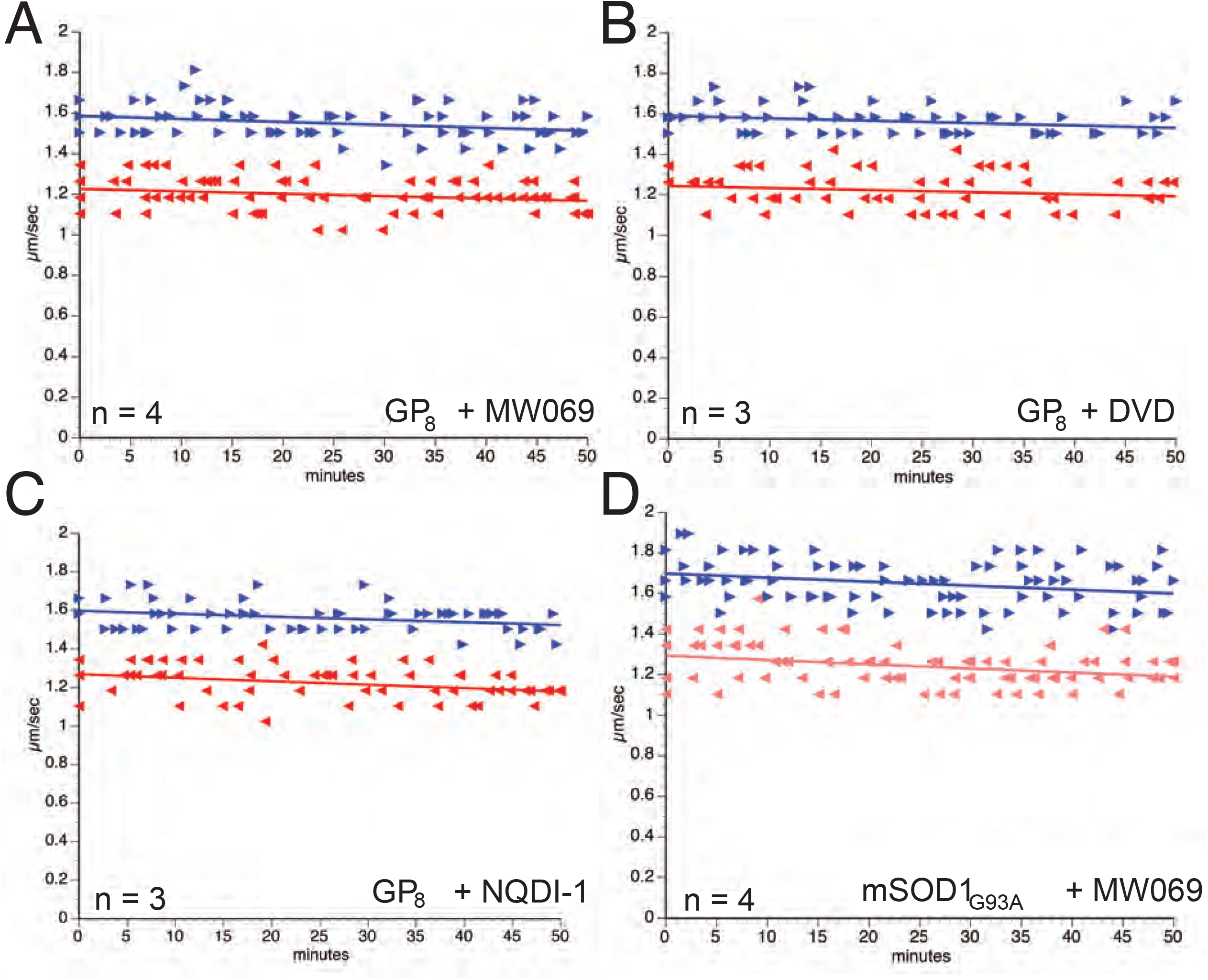
The inhibition of anterograde transport induced by GP(n_8_) peptide involves activation of kinases in the ASK1-p38 MAPK pathway. (**A**) MW01-2-069SRM (MW069), a selective inhibitor of p38α, blocked the inhibition of anterograde axonal transport induced by GP(n_8_) peptide repeat. (**B**) Co-perfusion of GP(n_8_) with 20µM DVD, a peptide inhibitor of a subset of MAPKKK, also prevented inhibition of anterograde transport. Interestingly, NQDI, an ASK1 inhibitor, had a similar protective effect **(C)**, suggesting that GP(n_8_)-induced toxicity on axonal transport is mediated by the ASK1-P38 pathway. Remarkably, our previous studies showed that MW069 (**D**), DVD peptide, and NQDI also prevented the inhibitory effect of mutant SOD1^[28, 29]^.

To provide additional evidence that inhibition of axonal transport by GP_8_ was through specific activation of P38 MAPK pathway, we performed metabolic labeling experiments to examine the phosphorylation of kinesin and various candidate kinases in axoplasms perfused with DPRs, as described previously^[45]^ (Fig 3). In the presence of ^32^P-γ-ATP, GP_8_ perfusion increased phosphorylation of Kinesin Heavy Chain (KHC) subunits, as shown before with mutant SOD1^[28]^ (Fig 3A-B). Immunoblot experiments further revealed increased levels of active (phosphorylated) P38 MAPKs in axoplasms perfused with GP, whereas levels of active GSK3 and ERK kinases remained unchanged (Fig 3C-F). Unlike GP_8_, GA_8_ perfusion had no effect on phospho-P38 levels (Fig 3G-H). Collectively, these results suggested that GP DPRs activate p38 MAPK and aberrant phosphorylation of the motor protein kinesin in the axonal compartment.

**Figure 3.**
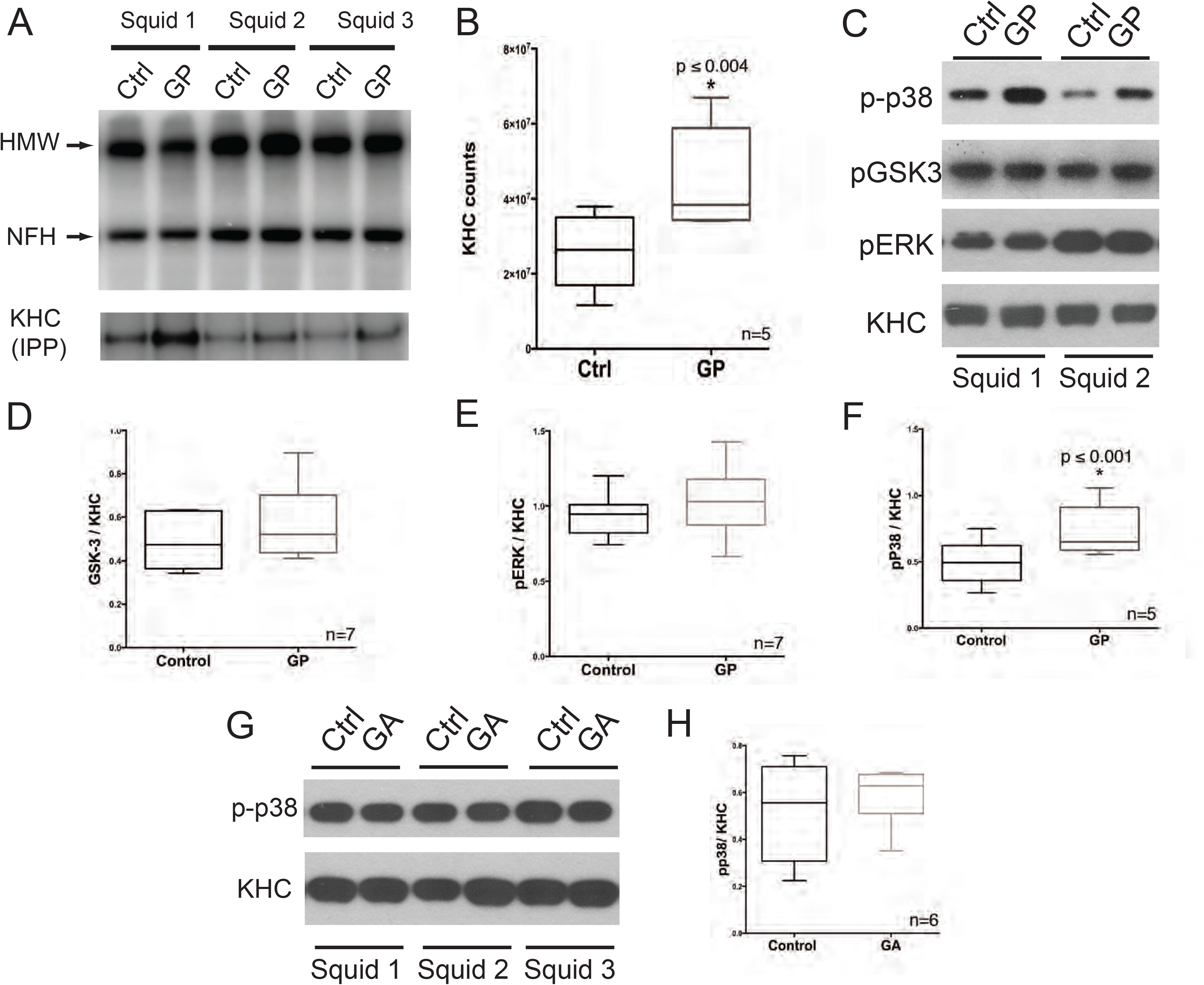
GP(n_8_) peptide increases kinesin phosphorylation *via* axon-autonomous activation of p38 kinases. Sister axons were extruded and perfused with control buffer (X/2, Ctrl) or with X/2 buffe plus 1µM GP(n_8_) dipeptide repeat (GP) in the presence of ^32^P-γ-ATP. After 50 min incubation, an aliquot of lysates was separated by SDS-PAGE and analyzed by autoradiography (**A**). High-Molecular-Weight Neurofilament (HMW) and Neurofilament Heavy Chain (NFH), major axoplasmic phosphoproteins in squid, are indicated. Kinesin was immunoprecipitated from the reminder lysate using H2, (an antibody against Kinesin Heavy Chain). Immunoprecipitated were separated by SDS-PAGE and ^32^P incorporation quantified by Phosphorimager scanning (KHC IPP) (**B**). Immunoblotting analysis of axoplasms showed increased activation/phosphorylation of p38 (p-p38) in axoplasms perfused with GP (n_8_) dipeptide, compared to “sister’ axons perfused with control buffer (Ctrl) **(C)**. Anti-kinesin-1 (KHC) antibody served as a loading control for axoplasmic protein. Quantitation of blots revealed no significant differences in levels of active GSK3 (**D**) or ERK kinases (**E**) but a significant increase in p-p38 in axoplasms perfused with GP(n_8_) peptide, compared to sister axons perfused with control buffer (**F**). (**G-H)**: Perfusion of GA(n_8_) peptide did not activate P38, as indicated by comparable phospho-p38 levels in western blots.

### GP dipeptide repeat protein inhibited synaptic transmission via ASK1

That GP DPRs inhibited axonal transport by activating a MAP kinase cascade, similar to mutant SOD1 that also inhibited synaptic transmission^[30]^, raised the question of whether GP and other DPRs also affected synaptic transmission^[26, 27, 30]^. To address this question, we used the squid giant synapse preparation. This model mimics mammalian neuromuscular junctions, where synthetic DPRs were microinjected into the presynaptic terminal site, and membrane potentials were simultaneously recorded in both presynaptic and postsynaptic sites to evaluate the strength of synaptic transmission (Fig 4A). While presynaptic injection of GR_8_ did not affect postsynaptic membrane potential (PSP) for over 250min after injection (Fig 4B), GP_8_ gradually inhibited PSP to the extent that postsynaptic action potential was no longer elicited within 200min after injection (Fig 4C). NQDI, a small molecule inhibitor of ASK1, blocked the inhibitory effect of GP_8_ and maintained steady synaptic transmission, similar to the synapses injected with GR_8_, as indicated by the constant PSP slope quantified over 250min (Fig 4D).

**Figure 4.**
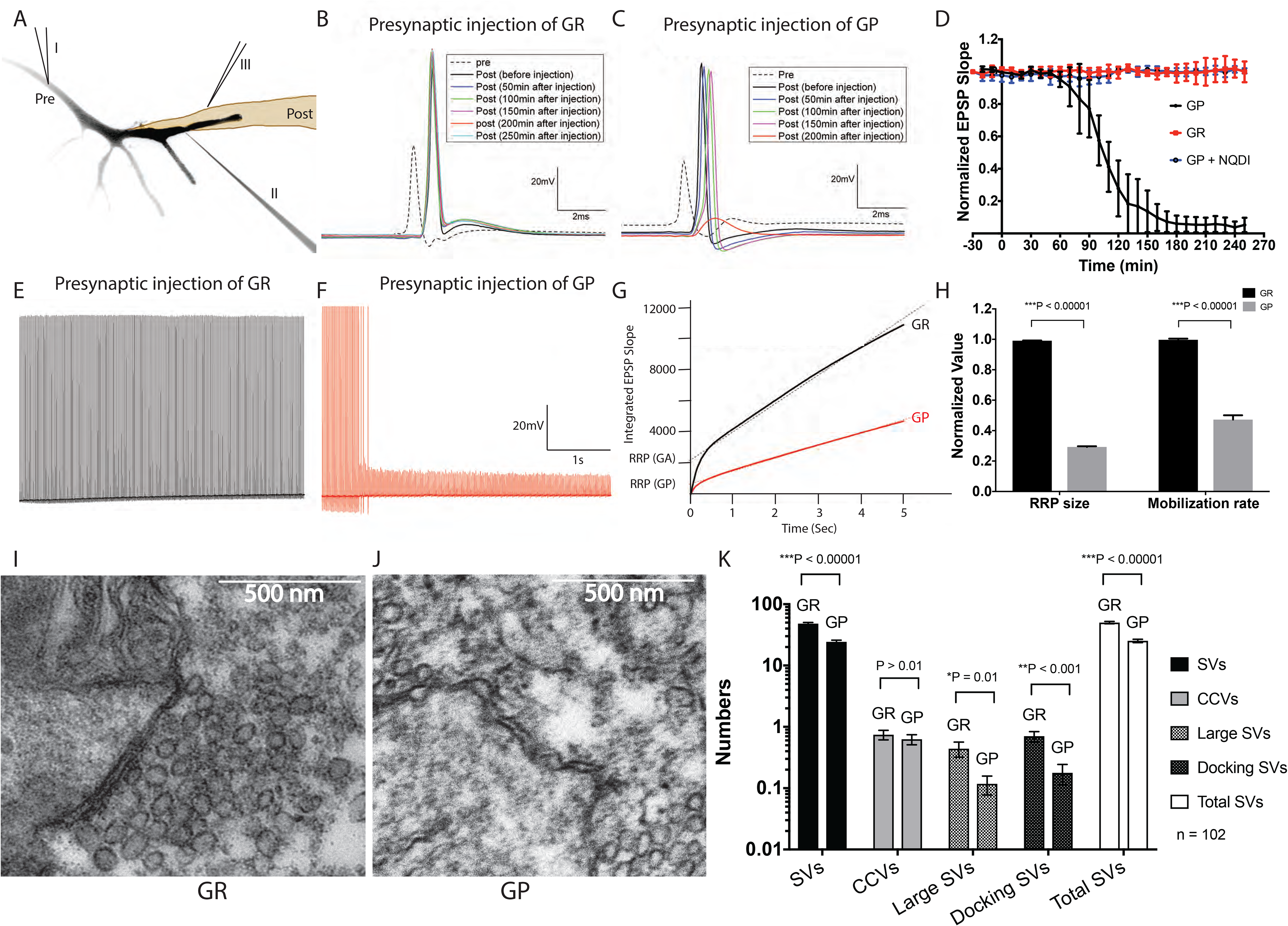
GP(n_8_) peptide inhibits synaptic transmission in the giant synapse. (**A**) Giant synapses were dissected and continuously superfused with oxygenated squid saline. GP(n_8_) or GR (n_8_) were injected into the presynaptic terminal (II) at 50 psi for 250ms/injection at 0.1Hz, and membrane potentials were recorded from both pre-(II) and post-synaptic (III) terminals. GR(n_8_) had no inhibition of synaptic transmission as the synapse kept firing for 250min **(B)**. In contrast, presynaptic injection of GP(n_8_) gradually reduced postsynaptic potential and eventually led to the failure in firing within 200min **(C)**. Quantitative plots **(D)** showed synaptic actions of GP(n_8_) and GR(n_8_) and suggested that NQDI, the same ASK1 inhibitor that prevented axonal transport defects in (Fig. 2C), also blocked the inhibitory effect of GP(n_8_) on synaptic transmission (n = 5). High-frequency stimulation at 50Hz for 5sec didn’t affect EPSP slope in GR-injected synapse **(E)** but significantly reduced PSP slope in GP(n_8_) injected synapse **(F)**. EPSP slopes during the 5sec train stimulation were integrated and plotted with the linear fit **(G**), suggesting reduced PRR size and mobilization rate of synaptic vesicles in the synapses injected with GP(n_8_), as compared to the ones injected with GR(n_8_) (n = 5) **(H)**. Electron Microscopy showed sufficient synaptic vesicles at the active zone of the GR(n_8_)-synapses **(I)** but a drastic decrease of available vesicles in GP(n_8_)-synapses **(J)**. Quantification of various classes of vesicles showed differences in total synaptic vesicles (SVs), large electron lucid vesicles, and docking vesicles, with similar numbers of clathrin-coated vesicles (CCV). Data were plotted from 100 synapses, injected with either GP(n_8_) or GR(n_8_) across 3 separate biological repeats (**K**).

Mutant SOD1 was previously shown to affect synaptic function by reducing available synaptic vesicle pools^[30]^. To examine whether GP_8_-induced synaptic defects also include an action on synaptic vesicle pool dynamics, trains of high-frequency stimulations (HFS, 50Hz for 5s, with 5s intervals between trains) were applied presynaptically 15min post-injection and postsynaptic depression of EPSPs were measured. Normally, a two-phase diminution of EPSP slope is observed across the 250 spikes^[30]^. The first phase is associated with the utilization of the Readily Releasable Pool (RRP) of synaptic vesicles, while the second phase reflects the Reserved Pool (RP). GR_8_-injected synapses quickly recovered from HFS (Fig 4F), suggesting abundant supplies of both RRP and RP. However, GP_8_-injected synapses were significantly inhibited (Fig 4E).

To further illustrate the vesicle pool dynamics, the EPSP slopes were integrated to generate a trace, where the last 50 time points were linearly fitted (Fig 4G). Both the relative size of the RRP, determined by the intersection of the linear fit with the y-axis, and the mobilization rate from the RP to RRP, determined by the slope of the linear fit, showed reductions in GP_8_-injected synapses, as compared with the GR_8_-injected synapses, which behave similarly to non-injected controls (Fig 4G-H). EM analysis of the active zones confirmed the reduction in total synaptic vesicles, including the large vesicles and docking vesicles, in synapses injected with GP_8_, but not in synapses injected with GR_8_ (Fig 4I-K). The effects of GP_8_ on synaptic physiology and morphology are similar to those observed following mutant G85R-SOD1 injection in the giant synapse^[30]^.

### C9ORF72-GP, PR, GR dipeptide repeat proteins induced axonal degeneration in mammalian neurons

To validate the results showing selected toxic effects of GP DPRs in the squid axon and synapse preparations in a mammalian system, we evaluated toxicity for all five C9-derived DPRs in primary cultured mammalian motor neurons. Transfection of motor neurons with recombinant mCherry constructs that had specific 25-mer DPRs added to their C-terminals showed that mCherry-GP_25_ produced axonal degeneration, while mCherry-GST control had no effect on motor neurons (Fig 5A-B). The effect on motor neurons was the same whether the mRNA sequence encoding for the mCherry-GP dipeptide used the GGGGCC hexanucleotide repeat or used various alternative codons to generate a dipeptide tail on the mCherry without nucleotide repeats at the RNA level (Fig 5C-D). The effects of mCherry-GP_25_ were mitigated by treatment with the P38 MAP kinase inhibitor SB203580 (Fig S1) or with an ASK1 inhibitor (NQDI) (Figs S2-S3), consistent with the rescuing effects of P38 MAP kinase/ASK1 inhibition in the squid axoplasm and giant synapse models. Also, as seen in the squid models, neither mCherry-GA_25_ nor mCherry-PA_25_ affected motor neuron viability regardless of DPRs being encoded by either GGGGCC repeats or alternate codons. Contrasting with results in axoplasm and the giant synapse models, transfection of motor neurons with either mCherry-PR_25_ or mCherry-GR_25_ constructs was toxic to mammalian motor neurons (Fig 5). As with mCherry-GP, toxicity was seen whether GGGGCC repeats or alternative codons were used (5C-D). However, unlike GP, inhibitors of P38 MAP kinase or ASK1 failed to rescue neurons expressing mCherry-PR_25_ and mCherry-GR_25_ (Figs S1-S3), suggesting that the toxic effects elicited by mCherry-PR_25_ and mCherry-GR_25_ involve a pathogenic mechanism distinct from that triggered by mCherry-GP_25_.

**Figure 5.**
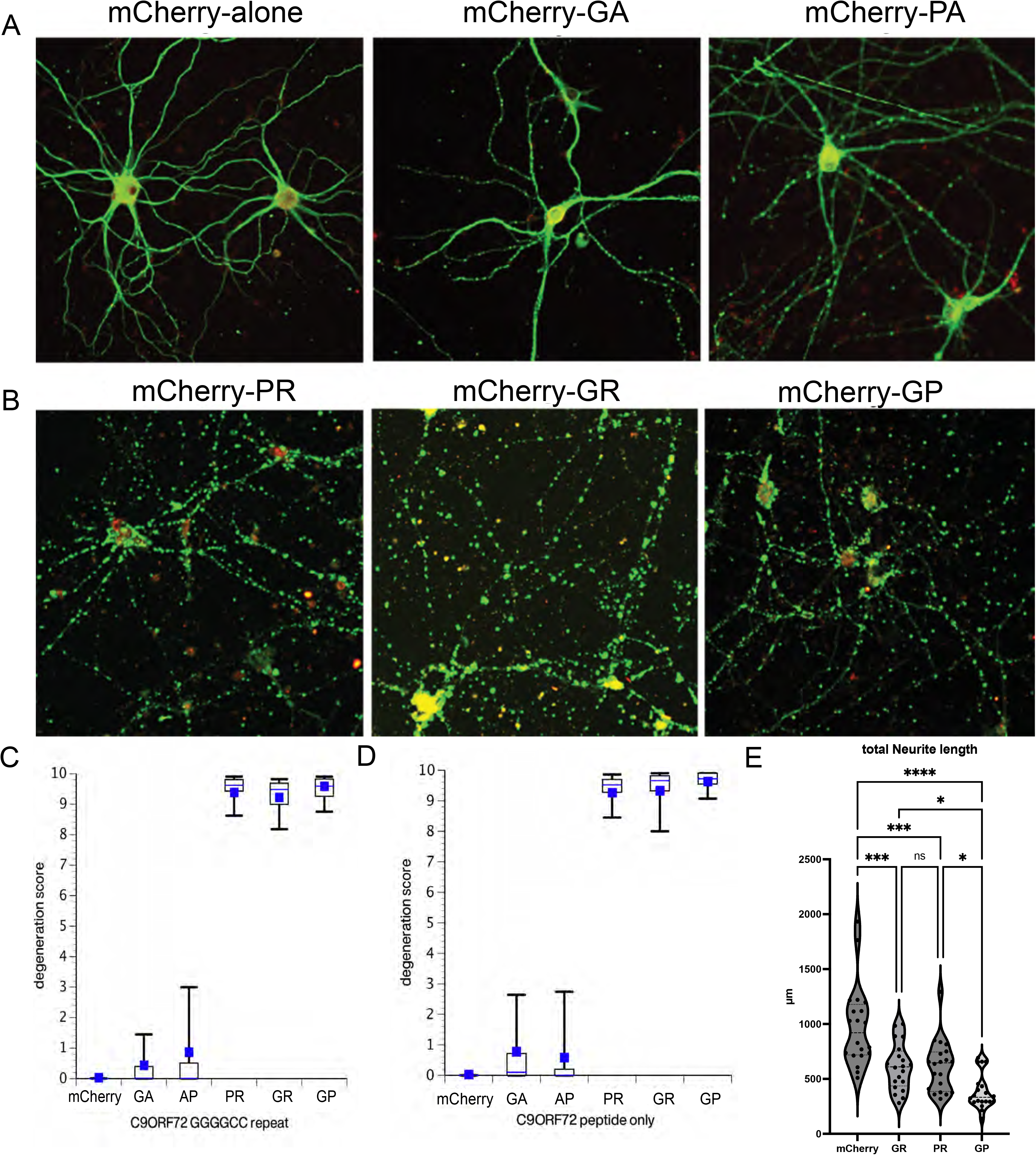
Toxicity of C9ORF72 Dipeptide Repeats in rat primary cultured motor neurons. Plasmids containing mCherry or mCherry fusion proteins with different dipeptide repeats (n=25 repeats) were used to transfect rat primary cultured spinal cord motor neurons. Neurons were transfected at 6DIV and fixed at 24h posttransfection. (**A**) Transfection with a plasmid expressing mCherry alone had no effect on neuronal viability. Similarly, transfection with mCherry-GA_25_ or -AP_25_ had little or no effect on neuronal morphology. (**B**) In contrast, motor neurons transfected with mCherry-PR_25_, -GR_25_ or -GP_25_ exhibited significant degeneration. This was in contrast to the effects of dipeptide repeats on axonal transport in axonal compartments or presynaptic terminals, where only GP dipeptides had significant effects. Quantitation of neuronal degeneration with each of the dipeptide repeats indicated that the toxicity was comparable whether the dipeptides were the result of adding GGGGCC hexanucleotide repeats to mCherry (**C**) or the dipeptides resulted from using alternate codons for the same dipeptides (**D**). This confirmed that the toxicity of all three toxic mCherry dipeptide repeats (-PR, -GR, and -GP) were the result of the dipeptide effects rather than RNA toxicity.

Although GP, GR, and PR DPRs were all toxic to cultured motor neurons, the phenotype of neurons expressing GP was distinct from that of neurons expressing either PR or GR (Figs 6, S4-S7). Specifically, motor neurons (12 DIV) transfected with mCherry-GP_25_ exhibited axonal degeneration with fragmented axons, accumulation of kinesin, and axonal swellings 10 hours after transfection, similar to effects elicited by SOD1(G93A) and FUS (R521G) (Fig. S4). However, the cell bodies and nuclei remained relatively intact with dispersed chromatin (DNA stained with DAPI) and an intact nuclear envelope (labeled by Lamin A/C) (Fig S5). In contrast, axons of motor neurons transfected with either mCherry-GR_25_ or mCherry-PR_25_ remained largely intact 10 hours after transfection, and kinesin distribution was not altered (Fig. S4). Instead, transfection with mCherry-GR_25_ or -PR_25_ produced neuronal degeneration in the soma while initially sparing the axonal compartment. Both mCherry-GR_25_ and -PR_25_ expressing cells exhibited striking changes in nuclear structure, with clumping of chromatin (DNA) and disruption of the nuclear envelope (Lamin A/C) (Figs S6-S7). The effect of transfecting neuron with an empty mCherry vector was minimal after 24 hours. While GP mainly affected axons causing “dying back” degeneration (i.e., from axon to cell body), PR and GR primarily affected nuclear structure consistent with a “dying forward” degeneration (i.e., from cell body to axon).

**Figure 6.**
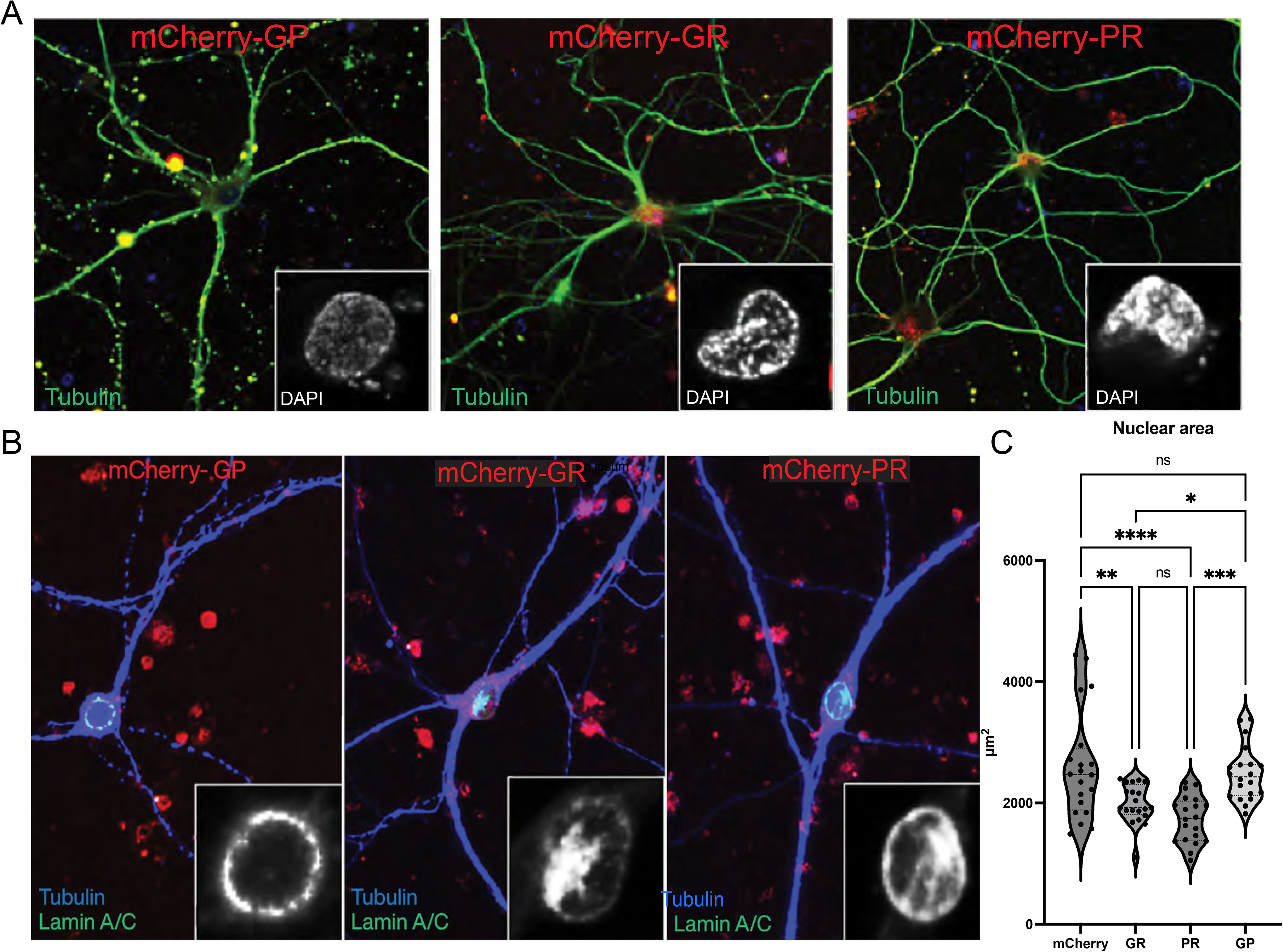
The GP dipeptides produce axonopathy, while PR and GR dipeptides exhibit nuclear changes. The mCherry-GP affects fast axonal transport, leading to a dying-back axonopathy. The nuclear envelope is intact (Lamin A/C) and chromatin is diffuse (DAPI). Axons are fragmented and truncated. In contrast, axons of neurons transfected with either mCherry-GR or mCherry-PR remain intact, but the chromatin (DAPI) is clumped and the nuclear envelope is breaking down (Lamin A/C). The nuclear area was quantified in (C), showing reductions in neurons expressing GR and PR but not GP. These different phenotypes indicate that there are at least two pathogenic mechanisms involved. Toxicity due to C9-GP dipeptides affect axonal transport and synaptic transmission through a pathway involving ASK1/P38 MAP kinase. In contrast, C9-GR and C9-PR dipeptides damage nuclear structure without affecting fast axonal transport or synaptic function, and their effects are not rescued by inhibitors of P38 or ASK1 (see supplemental figures 1-3).

In addition, mCherry-PR_25_ and mCherry-GR_25_ exhibited cytotoxicity for both neuronal and non-neuronal cells, whereas mCherry-GP_25_ was more toxic for motor neurons than non-neuronal cells. This difference was also observed when non-neuronal mammalian cells (COS 293) were transfected with the same DPR constructs (Figs S8-9). The mCherry-GP transfected cells were not significantly affected, with little evidence of toxicity. Cells transfected with mCherry-PR_25_ or mCherry-GR_25_ exhibited clear cytotoxicity with nuclear pathology that included aggregation of chromatin (DNA, Fig S8) and disruption of the nuclear envelope (Lamin A/C, Fig S9), similar to the effects of these dipeptides in neurons.

### Motor neurons from C9-ORF72 ALS patients showed a close association of C9ORF72-GP and active P38 kinase, both of which were significantly upregulated compared with controls

Finally, we performed immunohistochemical experiments in postmortem tissues from C9-ORF72 ALS and control patients to examine the potential relationships between C9-DPR levels and related signaling pathways identified in squid and primary culture neuron systems. Interestingly, we observed co-localization of C9-GP and active, phosphorylated P38 (pP38) in motor neurons undergoing degeneration, reported by reduced immunoreactivity of the neuronal-specific protein Tubulin Beta 3 Class III (Tubb3). Both GP and pP38 levels were higher in C9 motor neurons than in controls in both motor cortex (Fig 7) and spinal cord tissues (Fig S10). Structured Illumination Microscopy (SIM) with enhanced resolution confirmed the colocalization of GP and pP38 in both the nucleus and cytoplasmic compartments. The fluorescence intensities derived from anti-GP and anti-pP38 antibodies seem to correlate inversely with the anti-Tubb3-derived signal, suggesting a correlation with neurodegeneration.

**Figure 7.**
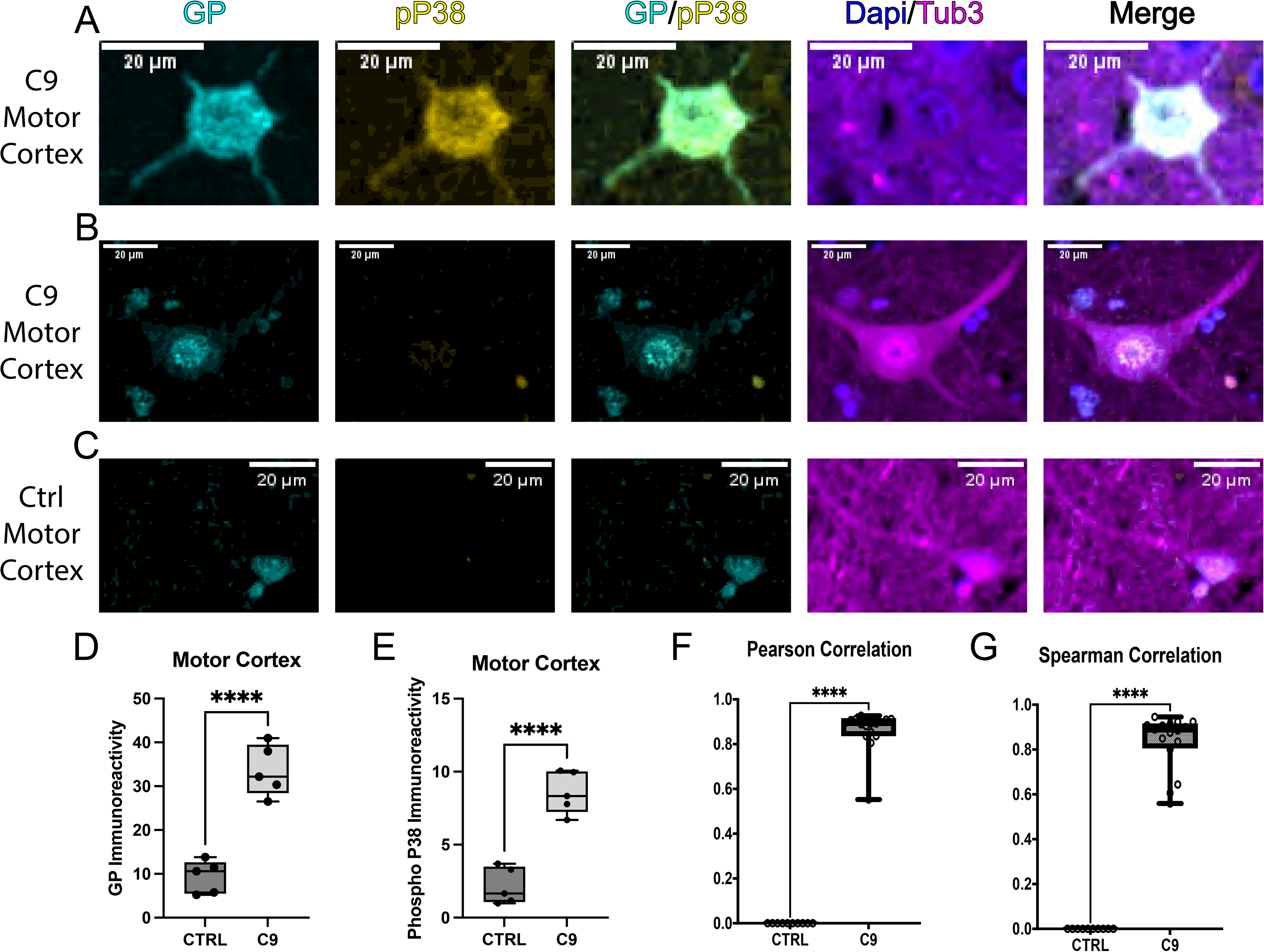
GP immunoreactivity is highly associated with P38 activity in the motor cortex of C9ORF72 patients. **(A)** GP and active P38 (phospho P38, pP38) were in close proximity in motor neurons, which showed reduced Tubb3 staining, indicating degeneration in C9 motor cortex. The immunoreactivities of GP and pP38 were in both cell body and processes. **(B)** A few surviving motor neurons showing normal Tubb3 staining showed little GP signal limited to the nucleus and no P38 activity, similar to those in control patients **(C)**. All results were quantified in **(D-G)**, n = 5/ group. This was further confirmed in the spinal cord using enhanced resolution microscopy (see supplemental figure 10).

Interestingly, cortical neurons from the superior frontal cortex of the C9-ORF72 FTD patients showed various degrees of nuclear envelope abnormalities, which were occasionally observed in aged control patients as well (Fig S11). In addition, some degenerating neurons also showed mislocalization of NUP107, a member of the nucleoporin family, in the cytoplasmic compartments. An anti-GR DPR antibody revealed GR aggregates close to the nuclear envelope in both nucleus and cytoplasm, though not co-localized with NUP107. While it is possible that misfolded GR directly interacts with NUPs, our data suggested that aberrant nucleocytoplasmic transport and mislocalization of NUPs may occur in C9ORF72 cortical neurons, and this process may be associated with GR aggregates and contribute to the nuclear damage, leading to a “dying forward” mechanism.

## DISCUSSION

ALS is defined by a clinical presentation in which both upper and lower motor neurons are compromised. The reasons for the selective vulnerability of these two neuronal populations continue to be debated and lead to a second question: do all forms of ALS share a common pathogenic mechanism, or does ALS present a group of distinct diseases with a common clinical presentation? Given that >90% of ALS cases are idiopathic with no family history and that the remaining familial cases of ALS have been linked to mutations in more than 40 different, seemingly unrelated genes, answers to this question are critical in the search for therapeutic interventions.

The discovery that expansion of an unstable hexanucleotide repeat in the C9ORF72 gene was associated with a surprisingly large number of both familial and sporadic forms of ALS provided a new challenge as well as an opportunity to understand ALS pathogenesis. The situation is further complicated by the fact that the same hexanucleotide expansion in C9ORF72 can also produce FTD and/or ALS symptoms in affected patients^[2, 3, 7]^. The frequency of different clinical presentations is roughly 45% ALS, 27% FTD, and 28% ALS/FTD^[46, 47]^. Futher, all three disease phenotypes can be observed in a single family with the same mutation. A single mutation that can variably affect two distinct neuronal populations is unusual, particularly since some other mutant genes associated with ALS, like SOD1, are rarely if ever associated with FTD^[4]^. The heterogeneity of neurological phenotypes seen with mutant C9ORF72 raises two questions: First, do ALS cases due to C9ORF72 mutations involve the same molecular mechanisms as ALS cases associated with other gene mutations and/or sporadic forms of ALS? Second, are the affected neuronal cell types and associated clinical symptoms of ALS and FTD the result of a common shared mechanisms or different pathogenic mechanisms?

The C9ORF72 mutation differs from other gene mutations associated with familial ALS in at least two ways. First, the mutation is an expansion of a hexanucleotide (GGGGCC) repeat that may have hundreds of repeats. Second, the hexanucleotide repeat is in an intron of the gene that is not normally transcribed. Based on these observations, three possible mechanisms for C9ORF72 toxicity were initially proposed: 1) loss of C9ORF72 protein function, 2) mutant C9ORF72 RNA-induced toxicity, and 3) toxicity of DPRs translated from these RNAs. Any of these mechanisms, alone or in combination, should explain the heterogeneity of disease phenotypes (i.e., ALS, ALS/FTD, and FTD) seen in patients bearing C9ORF72 mutations. On their own, ALS and FTD involve the degeneration of two distinct neuronal populations. These two neuronal populations appear to be differentially vulnerable because some mutant genes produce only ALS or only FTD. The variable presentation of disease phenotypes with C9ORF72 mutations raises the possibility that multiple pathogenic mechanisms are involved.

Loss of function alone was largely eliminated as a significant contributor to the development of either ALS or FTD^[9]^ because deletion of the C9ORF72 gene fails to produce either neurological phenotype but instead results in a fatal autoimmune disease^[7]^. This leaves RNA toxicity and/or DPR toxicity as candidate mechanisms. These two mechanisms are not mutually exclusive, so we tested both possibilities, with a focus on DPRs, in this study.

RNA toxicity is dependent on the hexanucleotide repeat forming unique RNA structures that might disrupt RNA processing or nucleocytoplasmic transport. The presence of RNA foci in affected cells provided support for this mechanism, and the focus has been primarily on changes in nucleocytoplasmic trafficking because FUS and TDP43, two other genes associated with familial ALS, have a role in nucleocytoplasmic trafficking of RNA^[48]^. In addition, cytoplasmic aggregates of wild type TDP43 can be seen in many cases of ALS and FTD in the absence of mutations. However, the question remains as to how RNA toxicity due to hexanucleotide expansion could produce either pure ALS or pure FTD as well as ALS/FTD.

The third candidate mechanism proposes that one or more of the five DPR-containing polypeptides generated by RAN translation of mutant C9ORF72 are toxic^[17–19]^. RAN-translated DPRs have been detected in multiple neurological diseases with repeat expansions and are implicated in pathogenesis^[18, 19]^. Variable levels of the five DPRs resulting from C9ORF72 mutations have been reported in cell bodies, distal axons, and synaptic terminals of affected cells^[20, 21]^. These DPRs could also affect RNA processing or trafficking. Alternatively, the DPRs could affect neuronal signaling pathways important for the regulation of fast axonal transport, as seen with mutations in SOD1^[28, 29, 37]^ and FUS^[39]^, or synaptic transmission, as shown for mutant SOD1^[30]^.

The possibility that mutant C9ORF72-related DPRs might affect axonal and synaptic signaling is particularly intriguing because one pathological change shared by affected motor neurons in both familial and sporadic cases of ALS is increased activity of the ASK1-MAP kinase pathway^[32, 33, 37]^. Our previous studies showed that several seemingly unrelated mutant genes associated with familial ALS activate this signaling pathway. These include familial ALS-linked mutant forms of SOD1^[28–30, 37, 38]^, an oxidized, misfolded form of wild type SOD1^[37]^, and mutant forms of the proteins FUS^[39]^ and profilin 1 (unpublished observations). Aberrant activation of the ASK1-p38/JNK MAP kinase signaling pathway in neurons has a number of potential consequences, including alterations of gene transcription, inhibition of axonal transport^[28, 29]^, and impairment of synaptic transmission^[30]^. These findings raised the question of whether C9ORF72 mutations could also affect the ASK1 MAP kinase pathway or if a different pathogenic mechanism was involved.

To evaluate axonal and synaptic pathogenicity of DPRs, we tested the pathogenicity of DPR-proteins associated with RAN translation of C9ORF72 in two unique model systems: isolated axoplasm and the giant synapse. These models allow analysis of DPR toxic effects independent of any contributions associated with the nucleus or RNA. Recombinant proteins containing each of the DPRs were perfused into the isolated axoplasm or injected into the giant synapse of the squid. Both axonal transport and synaptic transmission were inhibited by GP-DPR but not by any of the other four C9ORF72 DPRs. In previous studies with SOD1^[28, 29, 37]^ and FUS^[39]^, inhibition of axonal transport and synaptic activity was dependent on the activation of the ASK1-dependent kinase pathways. Similarly, the inhibitory effect of GP-DPRs on axonal transport was blocked by an ASK1 inhibitor and a P38 inhibitor. In the squid giant synapse preparation, synaptic inhibition by GP-DPR was also prevented by the ASK1 inhibitor, but not the P38 inhibitor (**Fig 4** and unpublished observations). These results were consistent with prior experiments demonstrating toxic effects of other familial ALS-associated mutant proteins (e.g., mutant SOD1, FUS) and sporadic ALS-associated oxidized SOD1 protein on axonal transport, as well as those showing inhibitory effects of ALS-associated mutant SOD1 protein on synaptic transmission *via* ASK1 activation^[30, 49]^. Altogether, these findings suggested a shared molecular component (i.e., ASK1-MAPK pathway) underlying common cellular dysfunctions (i.e., axonal transport inhibition and synaptic transmission deficits) that may lead to axonal degeneration in ALS-affected neurons, regardless of the mutant genes in familial cases and posttranslational modifications in sporadic cases, presenting ASK1 and downstream MAPKs as potential drug targets.

To validate these discoveries in mammalian neurons as well as explore additional pathological mechanisms, we expressed each of the five C9ORF72 DPR-proteins in mouse primary motor neurons. As predicted, GP-DPR induced axonal damage causing dying-back degeneration, similar to the effects seen with mutant SOD1, and initially sparing the neuronal cell body. However, unlike the results seen in isolated axoplasm and the presynaptic terminal of squid that lack neuronal cell bodies, two additional C9ORF72 DPRs, GR-DPR and PR-DPR, were also neurotoxic. Unlike GP-DPR, the PR- and GR-DPRs produced somal defects consistent with a dying-forward degeneration while initially sparing the axons. Significantly, though not surprisingly, inhibition of ASK1 rescued neurons expressing GP-DPR but failed to prevent GR/PR-DPR-mediated somal damage and neurodegeneration, suggesting that a different molecular mechanism for toxicity is involved.

As in the squid axoplasm, PA- and GA-DPRs had no effect on neurons. Since all five of the DPR-proteins included the same hexanucleotide expansion in different reading frames to produce specific DPRs, the lack of toxicity for PA- and GA-DPRs would not be consistent with a role for RNA toxicity in the observed phenotypes for GP-, PR- or GR-DPRs. To confirm that the observed toxicity of the DPR effects was independent of the RNA sequence with the hexanucleotide expansion, we repeated the experiments using transfected recombinant proteins in which the DPR sequences were encoded by alternative codons with no repeats. The RNAs produced by these plasmids would not have the same RNA structure but would express the same DPR. The phenotypes were identical for a given DPR, whether it was produced by plasmids with the GGGGCC hexanucleotide repeats or by alternative codons. The phenotypes seen in this study were the result of the corresponding DPRs, not by RNA. This does not exclude the possibility of an additional toxic gain function for RNAs, but it demonstrates that the DPRs are responsible for at least two distinct toxic gains of function differentially affecting the cell body and axon/synapse. Our human data from postmortem tissues confirmed the tight association of GP and active, phosphorylated P38 in motor neurons from C9ORF72 ALS patients, as well as the potential interaction of GR aggregates with the nuclear envelope in superior frontal cortical neurons from C9ORF72 FTD patients. A detailed pathological analysis in a bigger cohort of patients accompanied by mechanistic studies using patient iPSC neurons is currently underway to further elucidate the divergent mechanisms of C9-DPRs and the differential effects of individual C9-DPs in humans.

To determine whether C9ORF72 DPRs affect non-neuronal cells and characterize the phenotype in greater detail, we expressed the same C9ORF72-DPRs in COS293 cells. PR/GR-DPRs cause cell loss and nuclear envelope damage, similar to effects in neurons, while GP-DPR has no effect on nuclear structure or chromatin and exhibited no cytoxicity in COS293 cells. This is consistent with published literature showing toxicity of PR/GR-DPR in non-neuronal cells ^[50]^.

This also raises the possibility of glial toxicity due to PR/GR-induced somal toxicity in ALS, which may affect both motor and frontotemporal cortical neurons.

Neurons are dependent over a lifetime on maintaining active fast axonal transport and synaptic transmission over long distances, which may be as much as a meter or more for motor neurons in an adult human. Similarly, carefully regulated nucleocytoplasmic transport and nuclear function are essential for somal health and RNA and protein homeostasis in mature neurons. The heterogeneous phenotypes that can be seen with hexanucleotide expansion may result from the synergistic toxicity of GP-DPR and PR/GR-DPRs, and perhaps also GA when aggregated. Since RAN translation involves random initiation, the relative levels of each DPR in various subcellular compartments of a given patient may be variable. The GP-DPRs would be expected to affect upper and lower motor neurons through an ASK1-MAPK pathway and produce a dying back axonopathy, much like mutant and misfolded SOD1, suggesting that the GP-DPRs may be sufficient to produce an ALS phenotype. Since PR- or GR-DPRs activate a different pathogenic mechanism that is cytotoxic, potentially disrupting nuclear structure and function in both neuronal and non-neuronal cells, the presence of these DPRs may exacerbate the ALS phenotype as well as affecting additional neuronal populations such as in FTD. PR- and GR-DPRs could also produce dying forward degeneration in neurons by affecting glial cells and vasculature. The ability of mutant C9ORF72 to activate at least two independent, selective DPR-mediated pathogenic mechanisms may explain the existence of multiple distinct neuropathological phenotypes, with one pathogenic pathway contributing to ALS and the other to FTD.

In sum, our study demonstrated the toxic effects of GP-DPRs in the axonal and synaptic compartments that were mediated by an ASK1-p38 MAPK pathway, leading to dying back degeneration of motor neurons in ALS. In addition, the PR- and GR-DPRs activate a different pathogenic mechanism, which impacts the functionality of the nuclear compartment and may synergize to render motor neurons selectively vulnerable in ALS as well as the cortical neurons affected in FTD. The nuclear toxicity of PR and GR on neuronal and non-neuronal cells such as glia may further contribute to the disease pathology and include aberrant activation of additional kinase pathways with crosstalk with the ASK1 MAPK signaling. The multiplex disease pathology associated with C9ORF72 may call for a precision-medicine approach and cocktail therapy targeting multiple molecular components in a given patient.

## Methods

### Preparation of Dipeptides

Dipeptides corresponding to the five identified DPRs produced by RAN translation of expanded C9ORF72 hexanucleotide repeats (16-mer) were chemically synthesized and sequenced by the UIC Research Resources Center. Stocks of GP(8), GR(8), GA(8), PR(8) and PA(8) DPRs were prepared as a 5 mM (GR) or 10 mM (GP) stock in 50 mM HEPES, ph7.4 or 5 mM DMSO (GA).

### Vesicle Motility assays in isolated axoplasm

Axoplasms were extruded from giant axons of the squid Loligo pealii (Marine Biological Laboratory) as described previously ^[24]^. Briefly, the two largest axons are removed from wild-caught squid, and associated tissue is removed. Once cleaned, the axon can be extruded onto a coverslip to leave an intact cylinder axonal cytoplasm (axoplasm), which continues to maintain organelle trafficking in both directions for hours^[53, 54]^. Anterograde and retrograde transport is visualized using video-enhanced contrast differential interference contrast microscopy Zeiss Axiomat with a 100X, 1.3 n.a. objective, and DIC optics. A Hamamatsu Argus 20 and Model 2400 CCD camera system were used for image processing and analysis. Organelle velocities were measured interactively with a Photonics Microscopy C2117 video manipulator (Hamamatsu). Purified proteins (i.e., SOD1), DPRs and kinase inhibitors were all diluted to 1µM final concentration into X/2 buffer (175 mM potassium aspartate, 65 mM taurine, 35 mM betaine, 25 mM glycine, 10 mM HEPES, 6.5 mM MgCl_2_, 5 mM EGTA, 1.5 mM CaCl_2_, 0.5 mM glucose, pH 7.2) supplemented with 5 mM ATP) and 25 µl added to perfusion chambers (Fig 1). Both anterograde and retrograde FAT were analyzed by video-enhanced microscopy over 50 minutes, as before^[24]^.

### Biochemical experiments in isolated axoplasm

Biochemistry was conducted as described previously^[45]^. Using two axoplasms from a single squid, one was perfused with control buffer and one was perfused with 1µM GP_8_ in Buffer X/2 + 5µM ATP for comparison. For experiments shown in Fig 3A-B, γ-^32^P-ATP was included in the perfusate. After a 50 min incubation, axoplasms were homogenized and processed for kinesin immunoprecipitation using the H2 antibody against kinesin heavy chain then processed for autoradiography as described previously^[45]^. Briefly, axoplasm lysates were precleared using a mixture of Protein G agarose beads (Pierce), and non-immune mouse IgG-conjugated Sepharose beads (Jackson Immunoresearch) for 1 hour at RT. After a short spin, supernatants were incubated with H2 antibody and Protein G beads for 4 hours.

H2-kinesin immunocomplexes were recovered by centrifugation (3000 g_max_ for 30 seconds), washed four times with 1 ml lysis buffer, once with 50mM HEPES pH 7.4, and resuspended in Laemmli buffer. For immunoblots in Fig 3C, axoplasm lysates were separated by SDS-PAGE on 4-12% Bis-Tris gels (NuPage minigels, Invitrogen), using MOPS Running Buffer (Invitrogen) and transferred to PVDF using Towbin buffer supplemented with 10% (v/v) methanol (90 minutes at 400mA using Hoeffer TE22 apparatus). Immunoblots were blocked with 1% (w/v) non-fat dried milk diluted in TBST (25 mM Tris pH 7.2, 2.68 mM KCl, 136.8 mM NaCl, 0,01% Tween-20, ph: 7.2). Sodium Orthovanadate (1 mM) and Sodium Fluoride (10 mM) was used in all incubation steps involving the use of phosphoantibodies, after correcting for pH. Membranes were incubated with primary antibodies overnight at 4° C in 1% IgG free-BSA (Jackson Immunoresearch), and washed four times with 0.1% Tween-20 in TBS. Primary antibody binding was detected with HRP-conjugated anti-mouse, anti-rabbit or anti-goat antibodies (Jackson Immunoresearch), and visualized by chemiluminescence (ECL, Amersham).

### Giant Synapse Electrophysiology

The giant squid synapse is formed between the terminal finger of second order axons of the pre-nerve and the third order axon of the last stellate nerve, which is the giant axon used for axonal transport studies as well as for classical voltage-clamp studies of ion channels^[55]^. Giant synapse was prepared as described ^[30]^ and superfused continuously with oxygenated squid saline (455 mM NaCl, 54 mM MgCl_2_, 11 mM CaCl_2_, 10 mM KCl, 3 mM NaHCO_3_, and 10 mM HEPES, pH 7.2) at 10–15°C. One microelectrode for injecting was inserted in the presynaptic axon to inject current at 0.033 Hz for basal stimulation and at 50 Hz for HFS (each pulse is 2 µA for 2 ms for basal stimulation and 2 µA for 1 ms for HFS), near the palm where the second order axon enters the ganglion to branch and form multiple synapses with postsynaptic axons. At the terminal of the most medial branch, a second microelectrode was inserted to presynaptically inject proteins and reagents of interest as well as recording the presynaptic membrane potentials. Finally, at the postsynaptic terminal of the giant synapse, a third microelectrode was inserted near the medial presynaptic digit to record the postsynaptic membrane potentials. Electrodes 1 and 3 were filled with 3 M KCl while the second microelectrode allowed the injection of 50 µM SOD1 proteins or reagents of interest in 100 mM KCl^[56]^ at 0.1 Hz (each injection was 50 psi for 250 ms). The injection efficiency of peptides was monitored by a fluorescence microscope, which ensures comparable levels of infused peptides. Both presynaptic potential and PSP were recorded using sharp microelectrodes with an Axoclamp-2A amplifier (Axon Instrument) and data analyzed using Labview (National Instruments) software (Yulong Li). Raw waveforms were used to calculate PSP slope using the same parameters for all experiments. For data acquired under HFS, PSP slopes were analyzed, integrated, plotted, and fitted linearly using the last 50 time points to derive the vesicle mobilization rate (slope of the linear fit) and the ready-releasable pool (RRP) size (intersection with the *y*-axis).

### Electron Microscopy of the giant synapse

After recordings, synapses were prepared and imaged as published ^[30]^. Briefly, synapses were fixed in 4% glutaraldehyde and 2% paraformaldehyde in 0.1 M cacodylate buffer for 12 h at 4°C, then washed and postfixed in 1% osmium tetroxide, followed by blocking impregnation with 2% uranyl acetate in 0.1 M sodium acetate, pH 5.0 for 24 h. After washing, dehydration was performed in ethanol with increasing concentrations, followed by propylene oxide. Resin and propylene infiltration was conducted before embedding in silicone molds in the oven at 62°C for 72 h. Sectioning and imaging were performed following standard EM protocols^[57, 58]^. Vesicle density at the active zones (AZs) was determined as the number of vesicles per square micrometer. All EM samples were numbered, processed, and analyzed under blinded conditions until the last step when data had to be pooled.

### Plasmid preparation

Plasmids encoding mCherry-tagged versions of the dipeptides were constructed on the pcDNA3-mCherry vector by insertion of either 25 GGGGCC repeats and adjusting the reading frame to produce specific DPR or DNA encoding the same DPR but using alternative codons so that there were no nucleotide repeats. All constructs were designed to produce 25 repeats of a specific dipeptide repeat following the mCherry. The constructs allow us to compare toxicity with hexanucleotide repeats to that with the same DPR but a different nucleotide sequence. This evaluates the relative contribution of RNA toxicity and dipeptide toxicity to the observed phenotypes.

### Primary culture of motor neurons

E15 Spinal motor neurons were isolated from Sprague-Dawley rat embryos (E15) according to published protocols^[59]^. Briefly, spinal cords were dissected, trypsinized, triturated and then separated on a 9% Optiprep centrifugation. Motor neurons were plated on laminin-coated slides. Primary culture spinal cord motor neurons transfected with mCherry-tagged versions of C9 repeats in Ibidi chambers. At 12 DIV, neurons were transfected using the TransIT-X2 reagent (Mirus #MIR6000). For transfection, DNA (0.5ug) was mixed with 1.5uL X2 in 10uL unsupplemented neurobasal medium and incubated for 15 min at room temperature, and then mixed with media in each culture compartment. Motor neurons were fixed 24hr post-transfection with 4% paraformaldehyde, washed 3x 5 min in PBS, autofluorescence quenched with 50 mM NH4Cl for 15 min, washed 3x 5 min in PBS, permeabilized with 0.1% Triton-X100 for 10 min, washed 3x 5 min in PBS, nonspecific binding blocked with 5% milk for 1hr, washed 2x with PBS, incubated overnight at 4 C with primary antibody (DM1a, 1:1000, Sigma T6199), washed 3x 5 min in PBS, incubated for 45 min at room temperature with secondary antibody (AF488, 1:800, Invitrogen # A11001), washed 3 x 5 min in PBS, and covered in an aqueous anti-fade mounting medium (Vectashield #H-1000) for storage at 4C until imaged (Zeiss LSM710, 25x).

### Neuronal imaging and quantification

Neuronal morphology was examined in fixed neurons transfected with plasmids encoding mCherry versions of C9 dipeptide repeats with or without drug treatment. Degeneration scores were conducted triple-blinded. Images were obtained by one individual from samples, showing neurons with isolated neurites. Twenty or more images were collected for each condition. The second individual placed boxes on all neurite segments in the image that were isolated. The length of neurites in these boxes were measured by a third person and the length showing fragmented, beaded or aggregated phenotypes compared to the total length of the neurite in the box were calculated. All three individuals were blinded as to the experimental condition when selecting images and neurites or while making measurements. Neuritic degeneration was scored as a degeneration score, which is the ratio (degenerating neurite length)/(total neurite length) times 10. Data is shown as a box plot summarizing data from ≥20 images for each condition. (Figs 6 and S1).

### Human tissue collection, immunohistochemistry, and imaging

Postmortem human brain tissues were fixed in formalin, embedded in paraffin and stored in the Massachusetts Alzheimer’s Disease Research Center at MGH, under the strict ethical and IRB regulations from the Neuropathology core. Tissue blocks of the spinal cord, superior frontal cortex, and posterior frontal cortex were sectioned at 7µm thickness and mounted on super frost (+) glass slides. Sections were then processed by Leica Bond Autostainer through steps of bake and dewax, antigen retrieval (with citrate buffer for 30min, 95°C), permeabilization (0.4% triton in TBST for 10min, RT) and blocking (10% goat serum and 3% BSA in TBST for 45min, RT). Slides were then treated with Trueblack (Biotium #23007) for 30sec to eliminate lipofuscin autofluorescence and reduce autofluorescence from other sources, followed by TBS rinses. Tissue samples were then incubated with primary antibodies diluted in a blocking buffer (see Table 1 for a list of antibodies) for 60min at RT. After washing in TBS, tissues were incubated with secondary antibodies for 60min at RT), washing (TBST). Finally, tissues were incubated with Tubb3-alexa-488 (or Alexa-647) directly conjugated antibody (60min, RT), followed by washes (TBS), Hoechst staining (1µg/ml, 5min, RT), TBS rinses, and mounting with ProLong^TM^ Glass Antifade Mountant (ThermoFisher P36980).

**TABLE 1:**
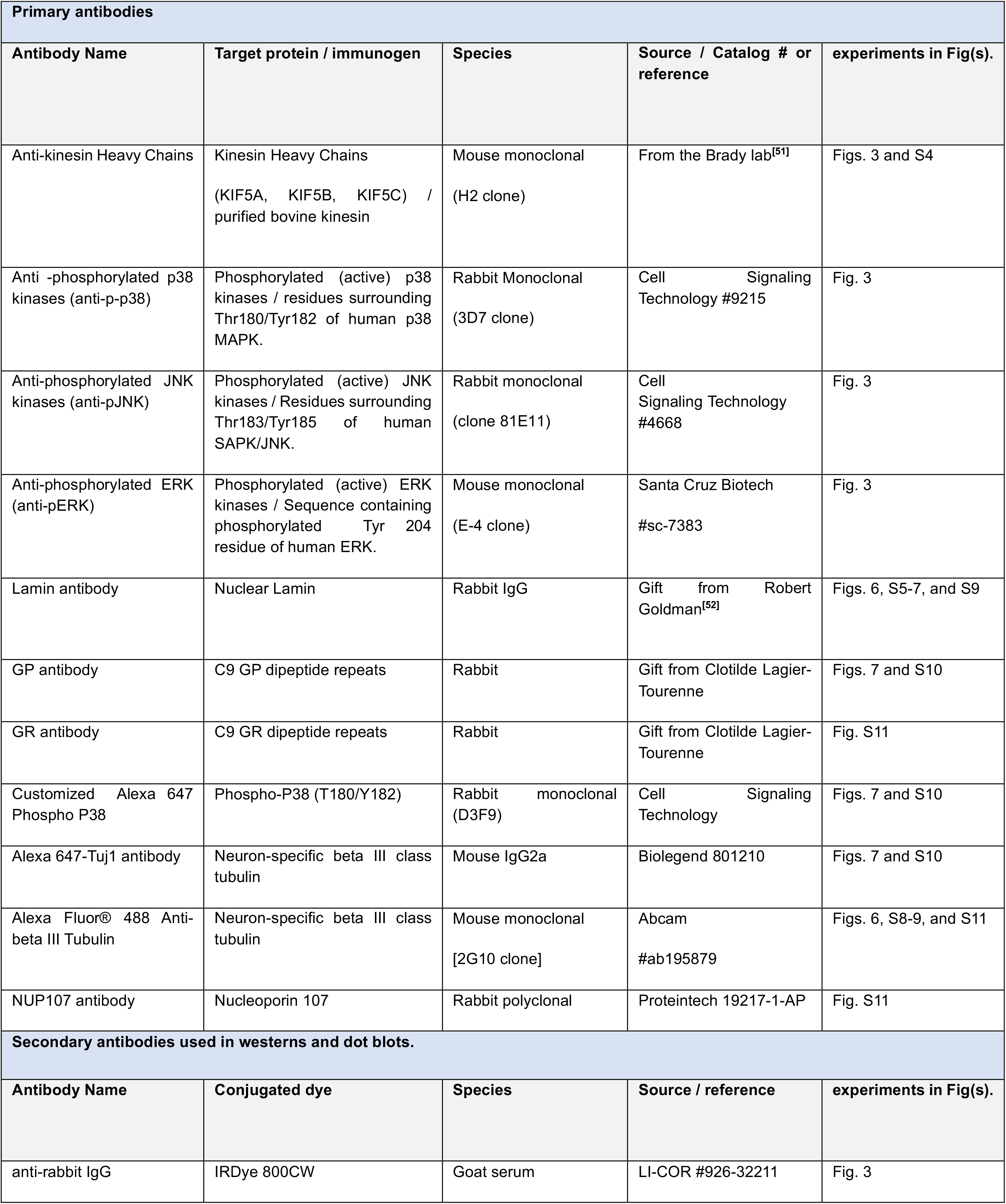
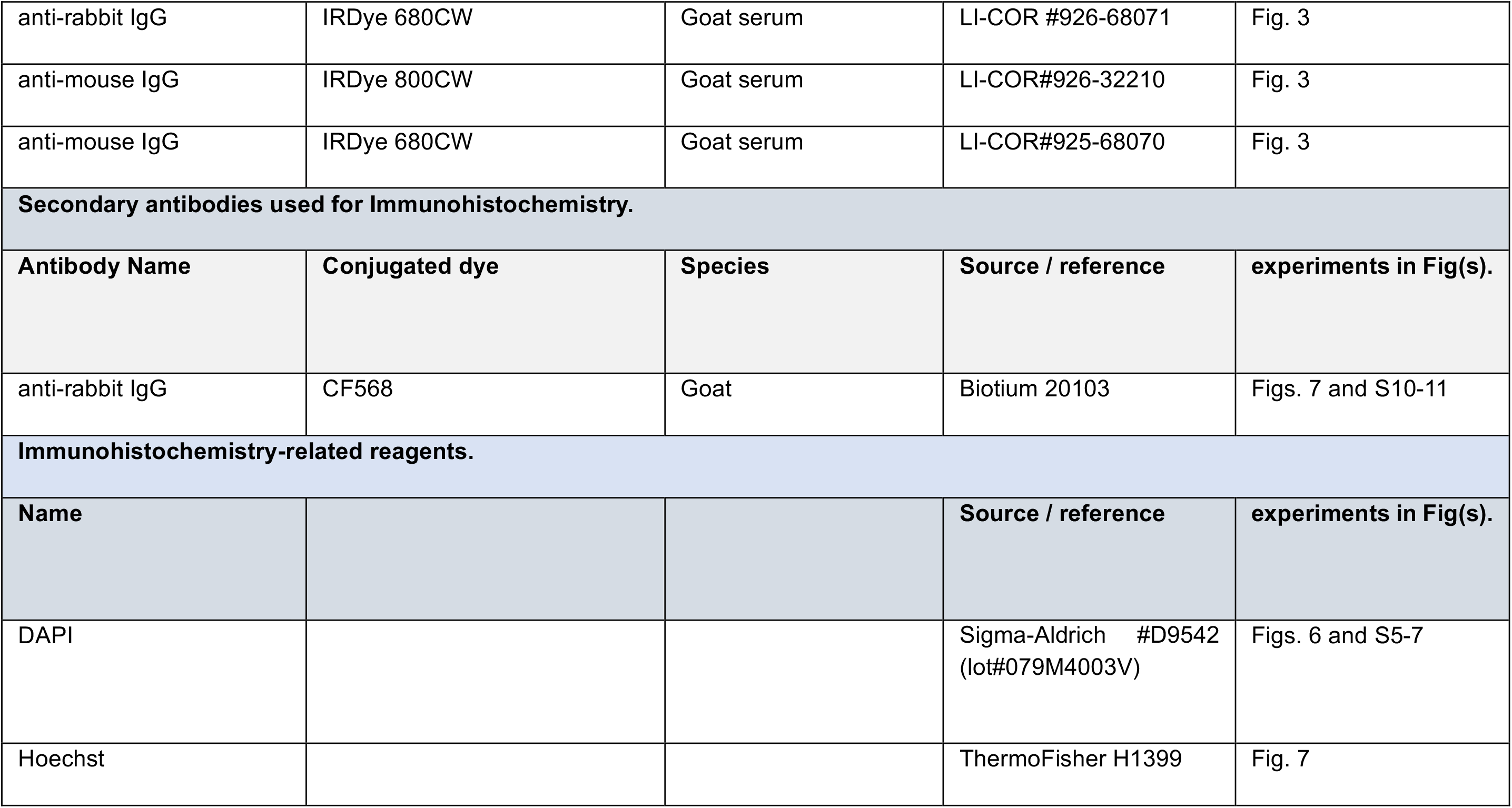
Antibodies used in this Study.

Epifluorescence imaging of the whole cortex or spinal cord section was performed automatically by Nanozoomer Slide Scanner (Hamamatsu Photonics) to avoid human bias in selecting regions of interest. All images were acquired using exactly the same parameters (e.g., exposure time) for all C9-ALS/FTD and control samples. Structured Illumination Microscopy was performed on Zeiss Elyra 7 and processed by SIM^2^ software (Zeiss) with alignment correction from calibration beads. Images were quantified by QuPath 0.4.3 for fluorescence intensity and by Fiji for colocalization by the user blinded to the sample genotype.

### Statistics

All experiments were repeated at least three times. The data were analyzed by one-way ANOVA followed by the Tukey *post hoc* test (or nonparametric multiple *t* tests, without assuming consistent SD) and plotted in Prism 7 (GraphPad software). Quantitative data were plotted as mean ± SEM, *p* values were calculated. For EM analysis, nested one-way ANOVA was performed.

## Author Contributions

YS, STB, and GM were responsible for the original conceptualization of the work and experimental design of all studies in this manuscript. YS and STB wrote the manuscript. SO, YS, GM, and STB performed all experiments in the squid axoplasm. YS, PC, and JEM performed all experiments in the squid giant synapse. MYT and BW performed all experiments in cell cultures. SO, PD, and YS performed all experiments in human tissues. NK contributed to biochemical experiments in squid axoplasm. SO, YS, MYT, BW, GM, and STB analyzed all the data. All authors reviewed and edited the final manuscript.

## Acknowledgments

We thank Dr. Marty Watterson (NWU) for the generous supply of p38 inhibitors, Dr. Clotilde Lagier-Tourenne for the kind gift of GP and GR antibodies, and Dr. Robert Goldman for the kind gift of Lamin antibody. We thank Drs. Merit Cudkowicz, Craig Blackstone, and Bradley Hyman for their full support of the Song Laboratory at MGH. The research was supported by NIH grants [P30AG062421 (Developmental Project) and AG072516], a Jack Satter Foundation grant, a Pape Adams ALS Transformative Scholar Award, Healey Center ALS Award, and an AARG grant from the Alzheimer’s Association to YS; NIH grants (R21NS120126 and R01NS118177) and a gift from Neurodegenerative Research Inc. to GM; and NIH grants (NS023868 and NS041170) to STB.

---

**Supplemental Figure S1. Inhibitors of P38 MAP kinase rescue mCherry-GP transfected neurons but not -PR or -GR transfected neurons. (A)** Degeneration of motor neurons transfected with mCherry-GP was rescued, at least partially, by treatment with the P38/JNK MAP kinase inhibitor SB203580. **(B)** Protection from -GP toxicity is mediated by inhibition of P38 MAP kinase as treatment with a selective inhibitor of P38a (193B) reduced degeneration due to -GP but did not reduce degeneration due to -PR and -GR. 193A, an inactive analogue of 193B didn’t prevent degeneration.

**Supplemental Figure S2. Live cell imaging of spinal motor neurons transfected with C9ORF72 dipeptide repeats.** Spinal motor neurons in culture were transfected with mCherry or mCherry-dipeptide, then stained with NeuO, a vital stain that preferentially labels neuronal membranes without affecting neuronal viability. Note that over 4 hours, neurons transfected with mCherry-GP cell body retained integrity, but the axons fragmented. In contrast, transfection with mCherry-PR or -GR leads to breakdown of the nucleus and clumping of DNA (insets stained with DAPI). Axons continue to be present.

**Supplemental Figure S3. Treatment of transfected neurons with NQDI-1, an ASK-1 inhibitor, rescues only -GP transfected neurons**. Over the course of 4 hours, mCherry-GP transfected neurons treated with NQDI-1 continued to grow and extend neurites. However, neurons transfected with either mCherry-PR or -GR continued to degenerate despite the presence of the ASK-1 inhibitor.

**Supplemental Figure S4. Transfection with GP, SOD1 (G93A), or Fus (R521G), but not with PR leads to kinesin accumulation and axonal damage.** mCherry-GP and two mutant gene products that cause familial ALS, mCherry-SOD1(G93A) and mCherry-Fus (R521G) all activate the ASK1/P38 MAP kinase signaling pathway and inhibit fast axonal transport by phosphorylation of kinesin. All three led to the appearance of kinesin aggregates in axons, reduction of overall axonal kinesin staining, and axonal pathology. In contrast, PR, which does not affect ASK1/P38 MAP kinase or axonal transport, showed kinesin staining of axons in speckles with little or no evidence of kinesin aggregates. Motor neurons were fixed 10 hours after transfection and processed for immunoflurorescence microscopy.

**Supplemental Figure S5. Nuclei in neurons transfected with mCherry-GP are normal.** Neurons transfected with mCherry-GP do not exhibit alterations in the nuclear envelope as visualized by the Lamin A/C antibody. Concurrently, DNA stained with DAPI is diffuse, consistent with normal chromatin structure.

**Supplemental Figure S6. Transfection with mCherry-GR affects nuclear structures.** Neurons transfected with mCherry-GR consistently show alterations in the nuclear envelope as visualized by the Lamin A/C antibody. Concurrently, DNA stained with DAPI also exhibits clumping.

**Supplemental Figure S7. Transfection with mCherry-PR affects nuclear structures.** Neurons transfected with mCherry-PR similarly show alterations in the nuclear envelope as visualized by the Lamin A/C antibody. Concurrently, DNA stained with DAPI also exhibits clumping.

**Supplemental Figure S8. Transfection with mCherry-PR or -GR is cytotoxic and exhibits DNA aggregates, but mCherry-GP transfected cells do not.** To evaluate the effects of C9ORF72 dipeptides on non-neuronal cells, COS 293 human embryonic kidney cells were transfected with mCherry-dipeptides. The -PR and -GR transfected cells exhibit DNA aggregates (DAPI) and disrupted microtubules, which indicate cytotoxicity. In contrast, the -GP transfected cells appear normal and viable.

**Supplemental Figure S9. Transfection with mCherry-PR or -GR is cytotoxic and transfected cells exhibit nuclear envelope alterations.** To evaluate the effects of C9ORF72 dipeptides on non-neuronal cells, COS 293 human embryonic kidney cells were transfected with mCherry-dipeptides. The -PR and -GR transfected cells exhibit nuclear envelope defects, as evidenced by lamin A/C-mediated invaginations in the nuclear membrane, which indicate nuclear toxicity. In contrast, the nuclei of -GP transfected cells appear normal with smooth and round nuclear envelopes.

**Supplemental Figure S10. Motor neurons in the spinal cord from C9ORF72-ALS patient showed increased immunoreactivity of GP and pP38 as well as their tight association by enhanced resolution SIM imaging. (A and B)** Two motor neurons with various levels of Tubb3 showed both cytoplasmic and nuclear GP and pP38 immunoreactivity in C9OGR72 patients. GP and pP38 were closely associated with each other. **(C and D)** Two motor neurons from control patients showed strong tubb3 signals with minimal GP and pP38 immunoreactivity.

**Supplemental Figure S11. Superior frontal cortical neurons from the C9ORF72-FTD patients exhibited aberrant nuclear structures and association with GR. (A)** Neurons at different stages showed various degrees of nuclear pore abnormality as labeled by NUP107. NUP107 completely mislocalized to the cytoplasm in one neuron, where the nuclear pore structure was lost. GR aggregates were associated with NUP107, though not co-localized, and may directly interact with nuclear pores to induce or worsen the structural changes of the nuclear envelope. However, nuclear damage may also be indirect of GR accumulation. This was more evident in the mortgage images of 90 Z-stacks by SIM imaging in **(B)**.

## Notes

### Competing Interest Statement

The authors have declared no competing interest.

### Summary of Updates

This revised manuscript has included new human data from C9ORF72 patients and their controls.

