## Supplemental Figures and Figure Legends for "Divergent Molecular Pathways for Toxicity of Selected Mutant C9ORF72-derived Dipeptide Repeats"

Figure S1

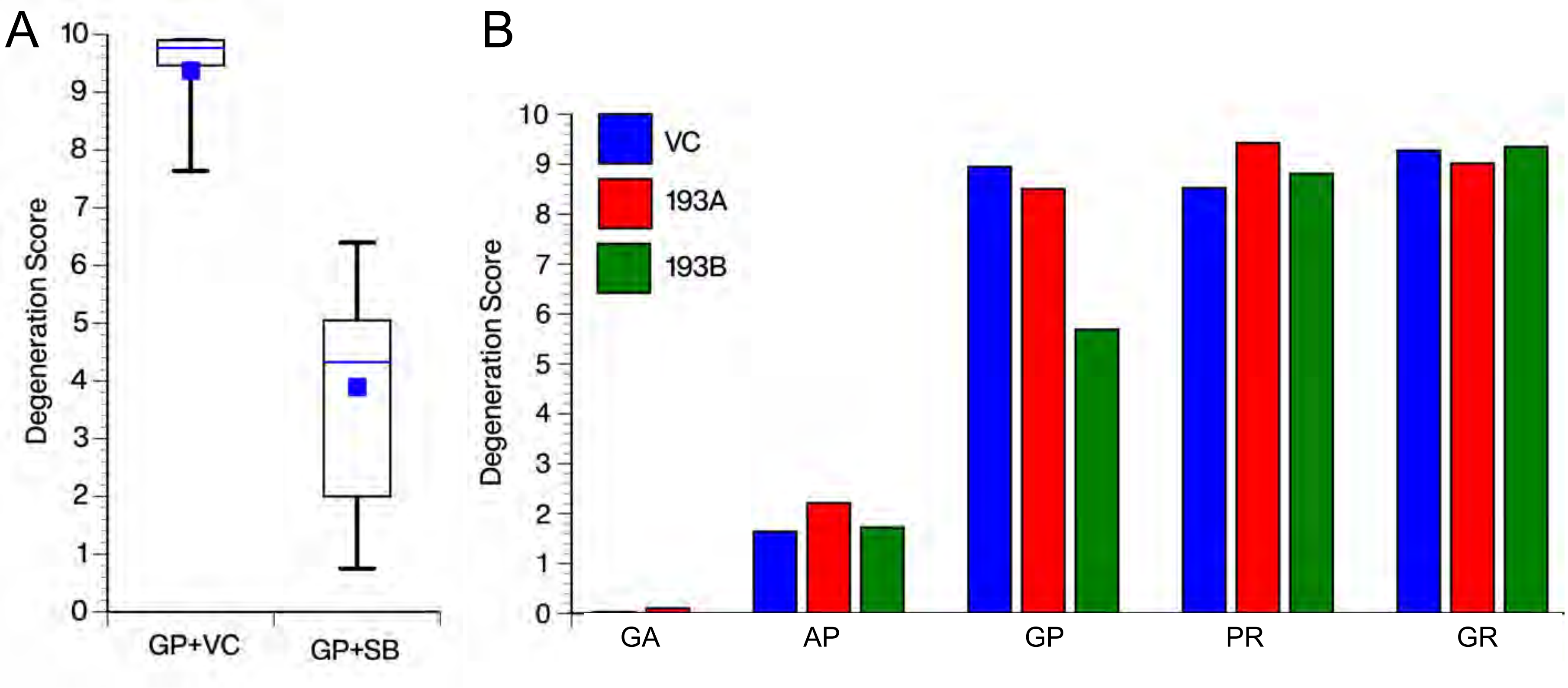
A
B
Degeneration Score
10
9
8
7
6
5
4
3
2
1
0
GP+VC
GP+SB
VC
193A
193B
GA
AP
GP
PR
GR

### Figure S2

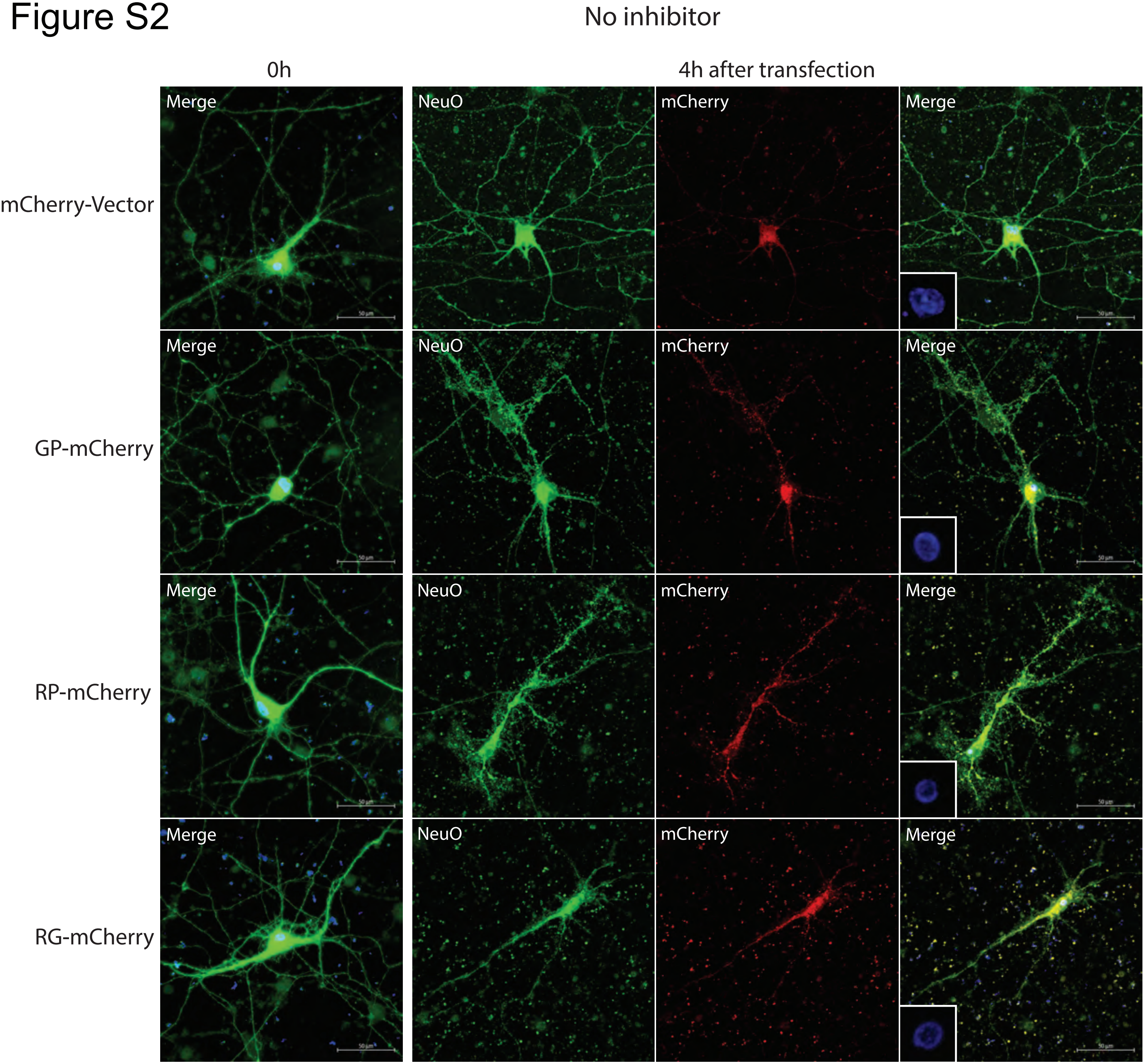
No inhibitor
0h
4h after transfection
mCherry-Vector
GP-mCherry
RP-mCherry
RG-mCherry
Merge
NeuO
mCherry
Merge

### Figure S3

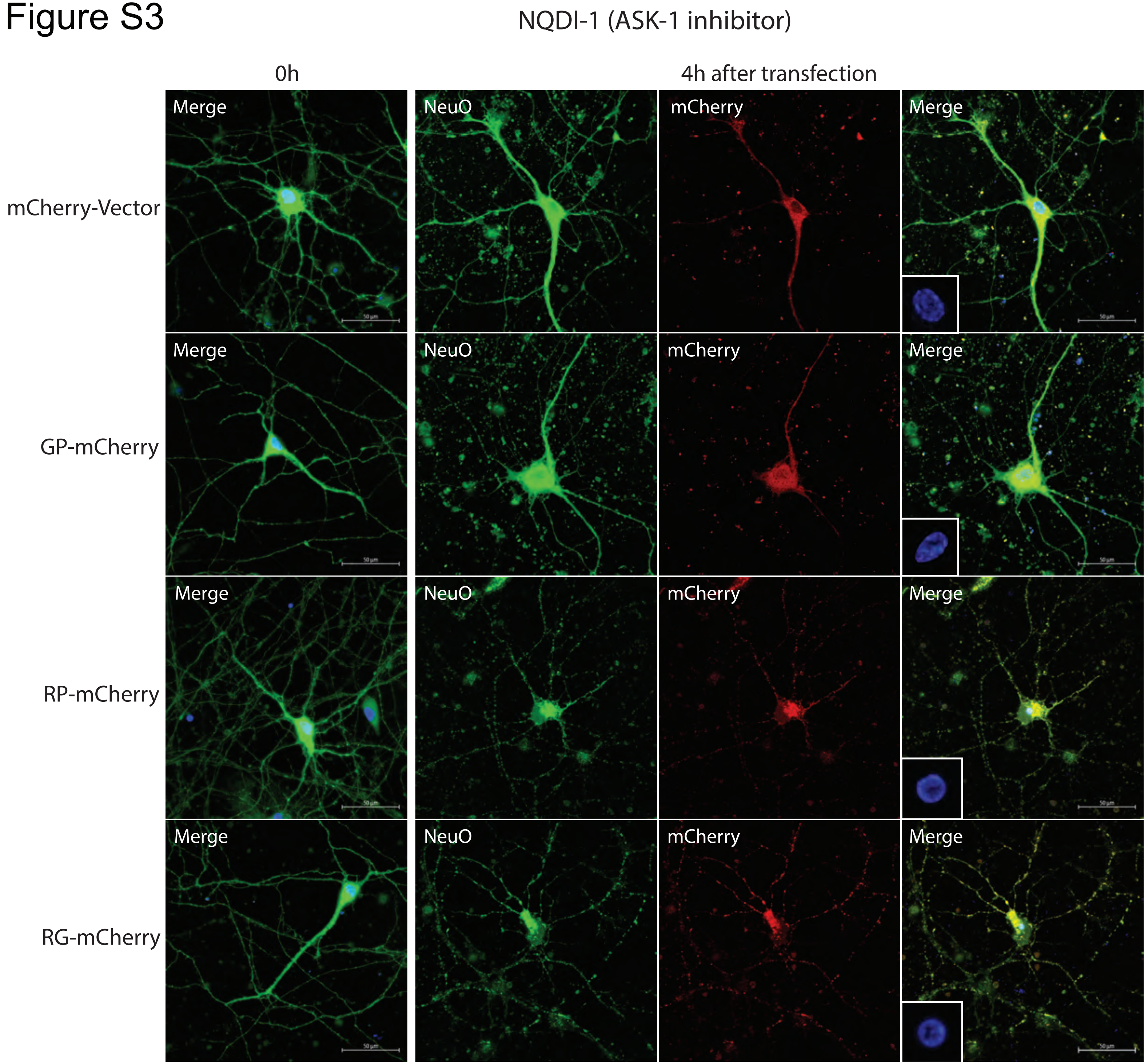
NQDI-1 (ASK-1 inhibitor)
0h
4h after transfection
mCherry-Vector
GP-mCherry
RP-mCherry
RG-mCherry
Merge
NeuO
mCherry
Merge

Figure S4

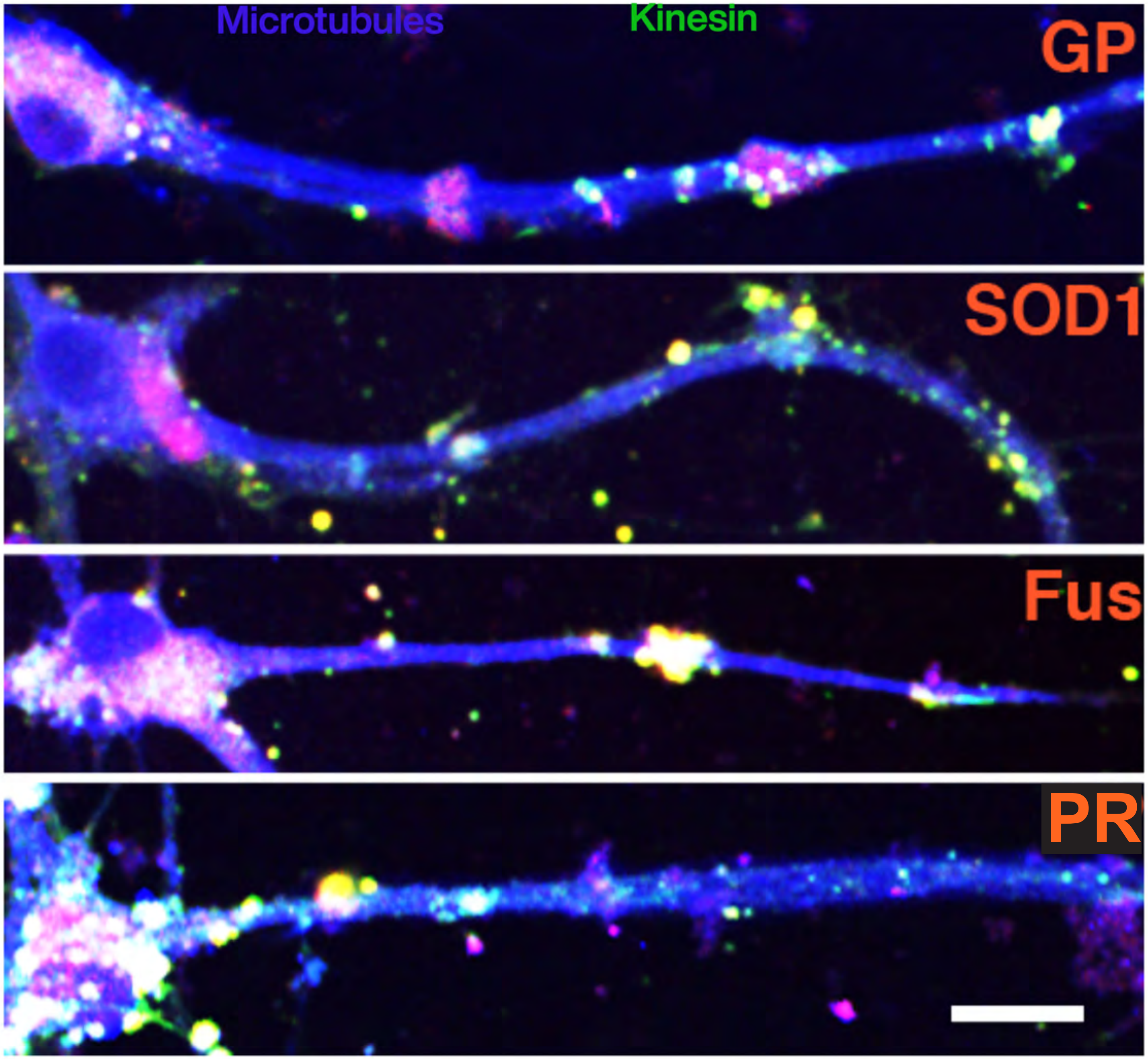

Figure S5

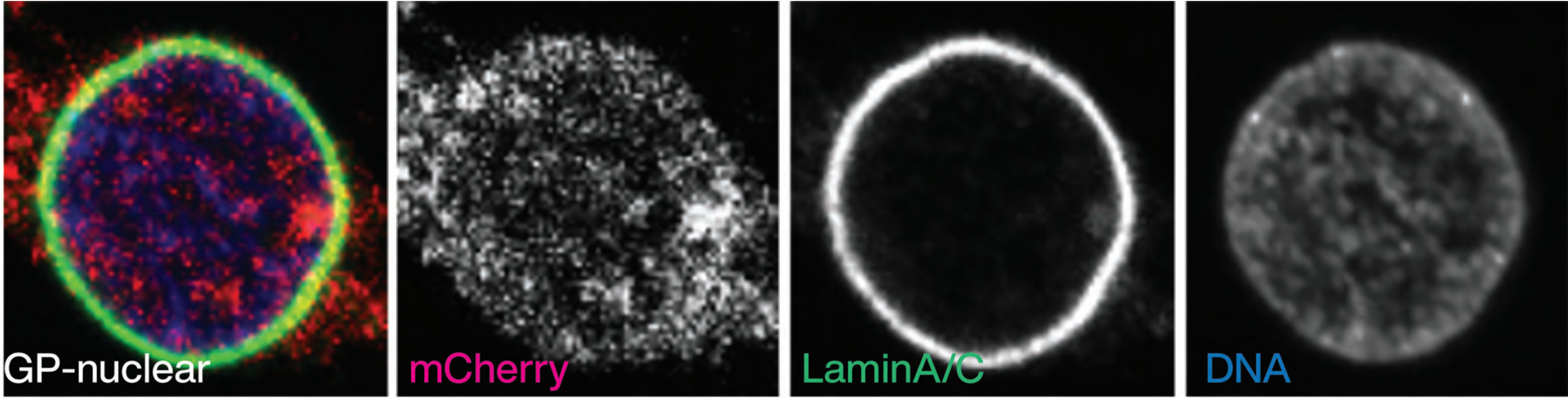

Figure S6

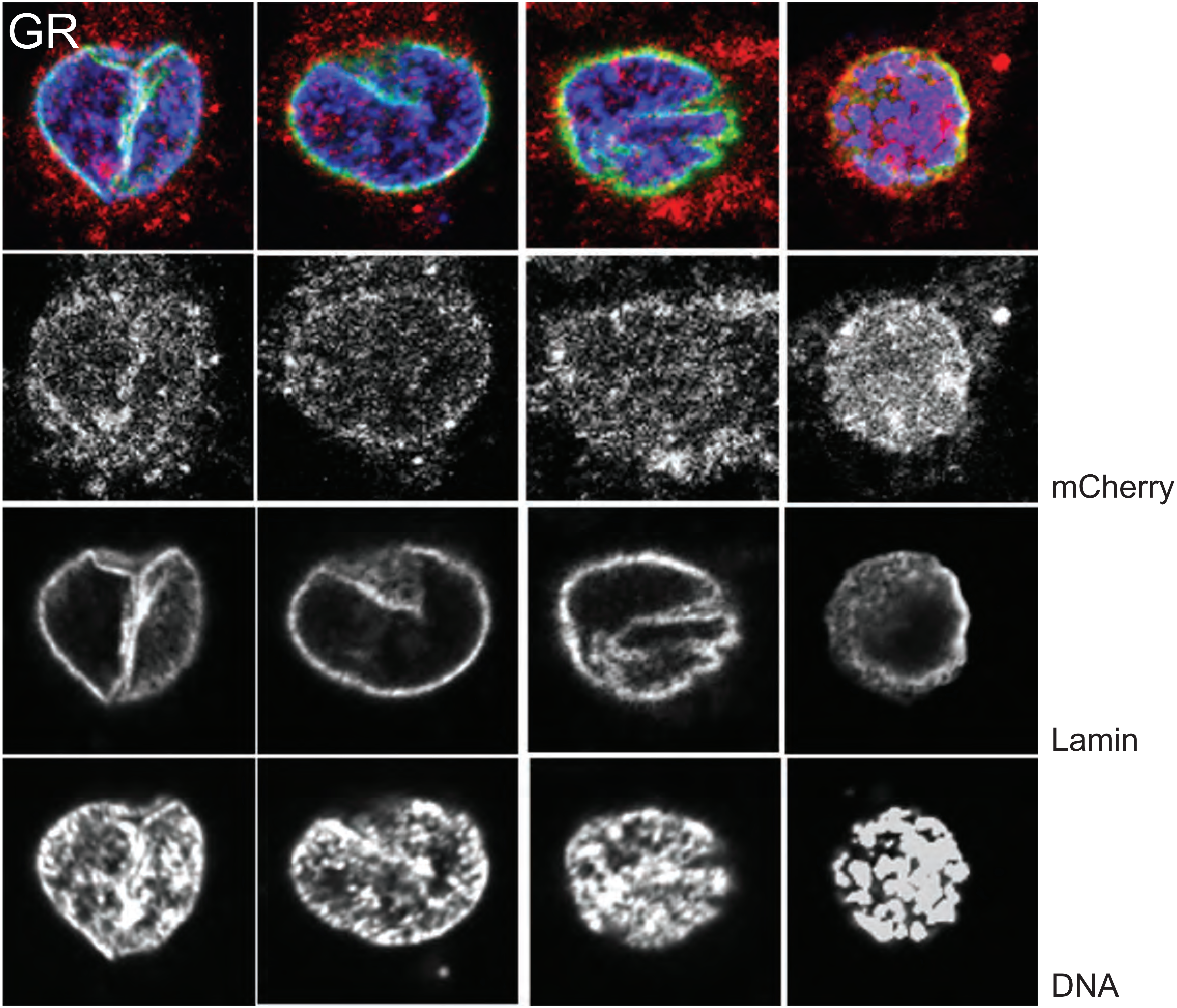

Figure S7

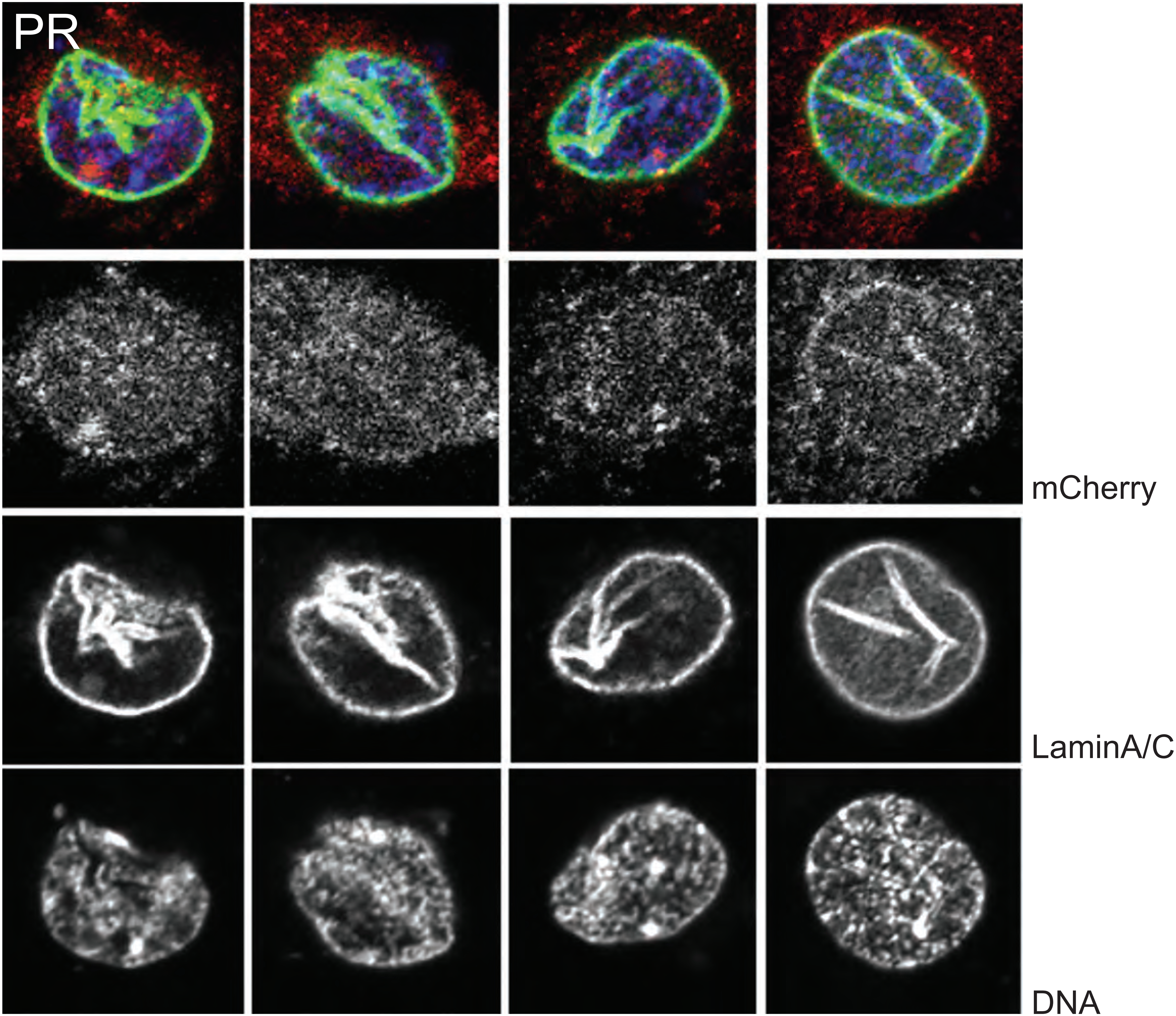

Figure S8

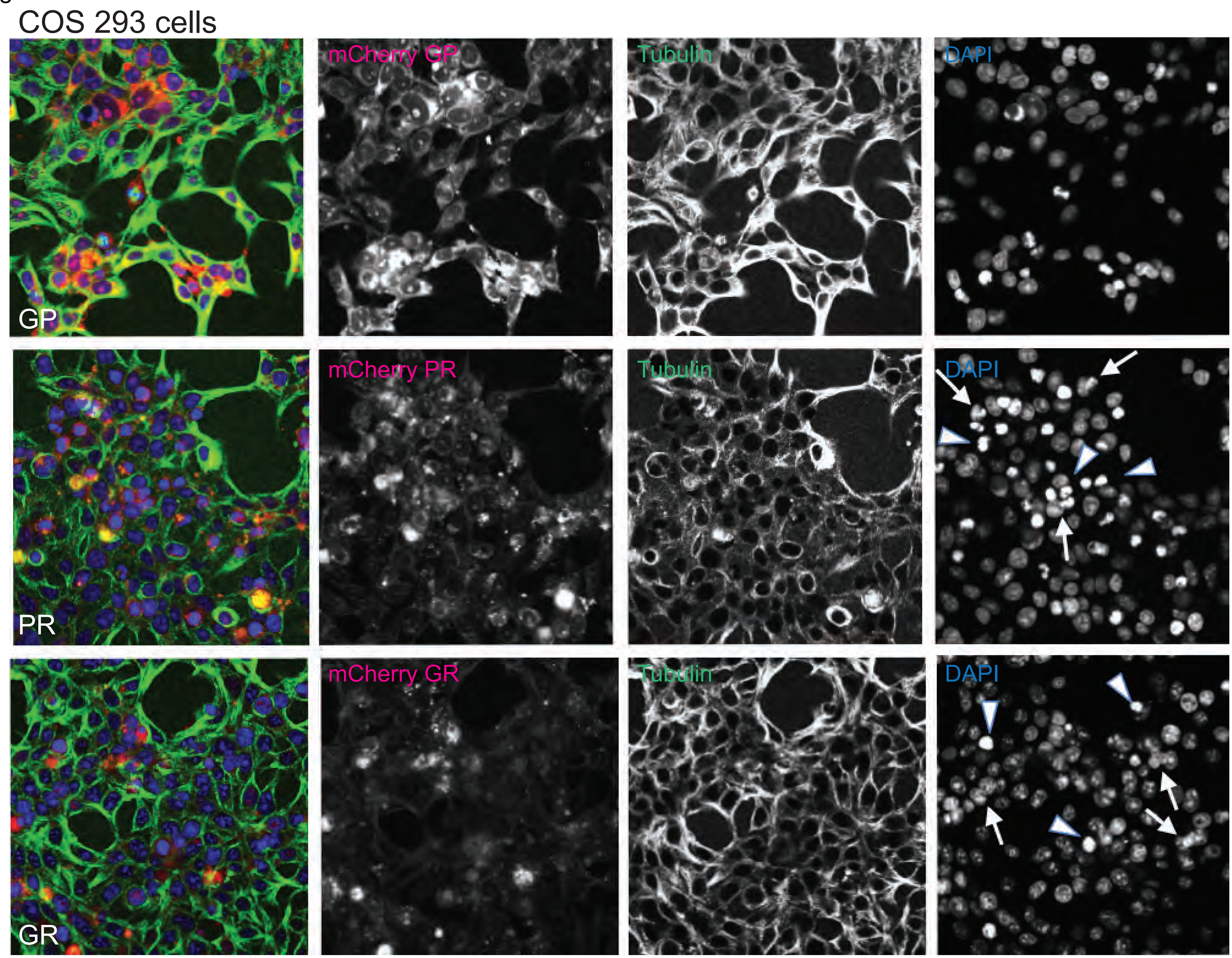
COS 293 cells
GP
mCherry GP
Tubulin
DAPI
PR
mCherry PR
Tubulin
DAPI
GR
mCherry GR
Tubulin
DAPI

Figure S9

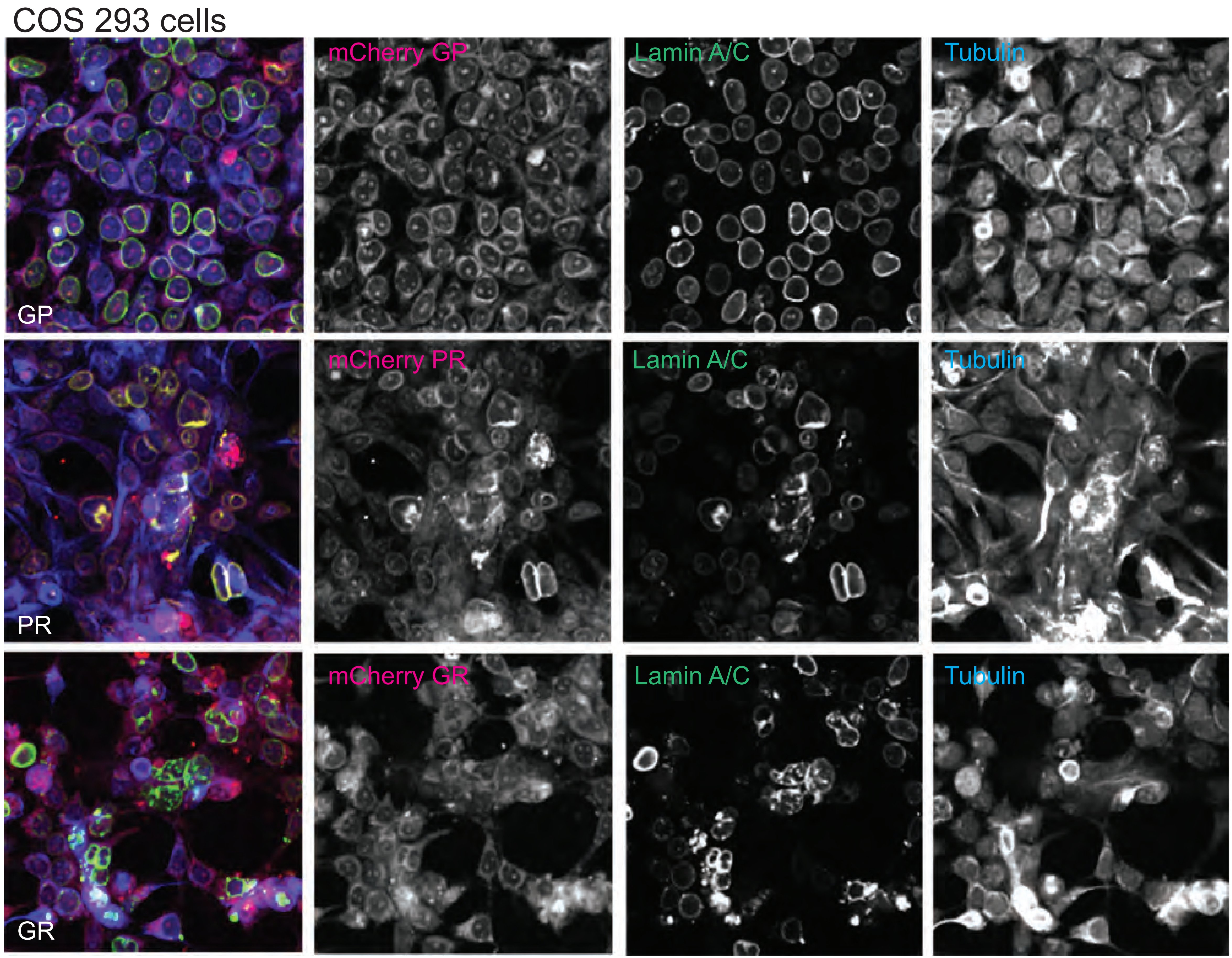
COS 293 cells
mCherry GP
Lamin A/C
Tubulin
GP
mCherry PR
Lamin A/C
Tubulin
PR
mCherry GR
Lamin A/C
Tubulin
GR

Figure S10

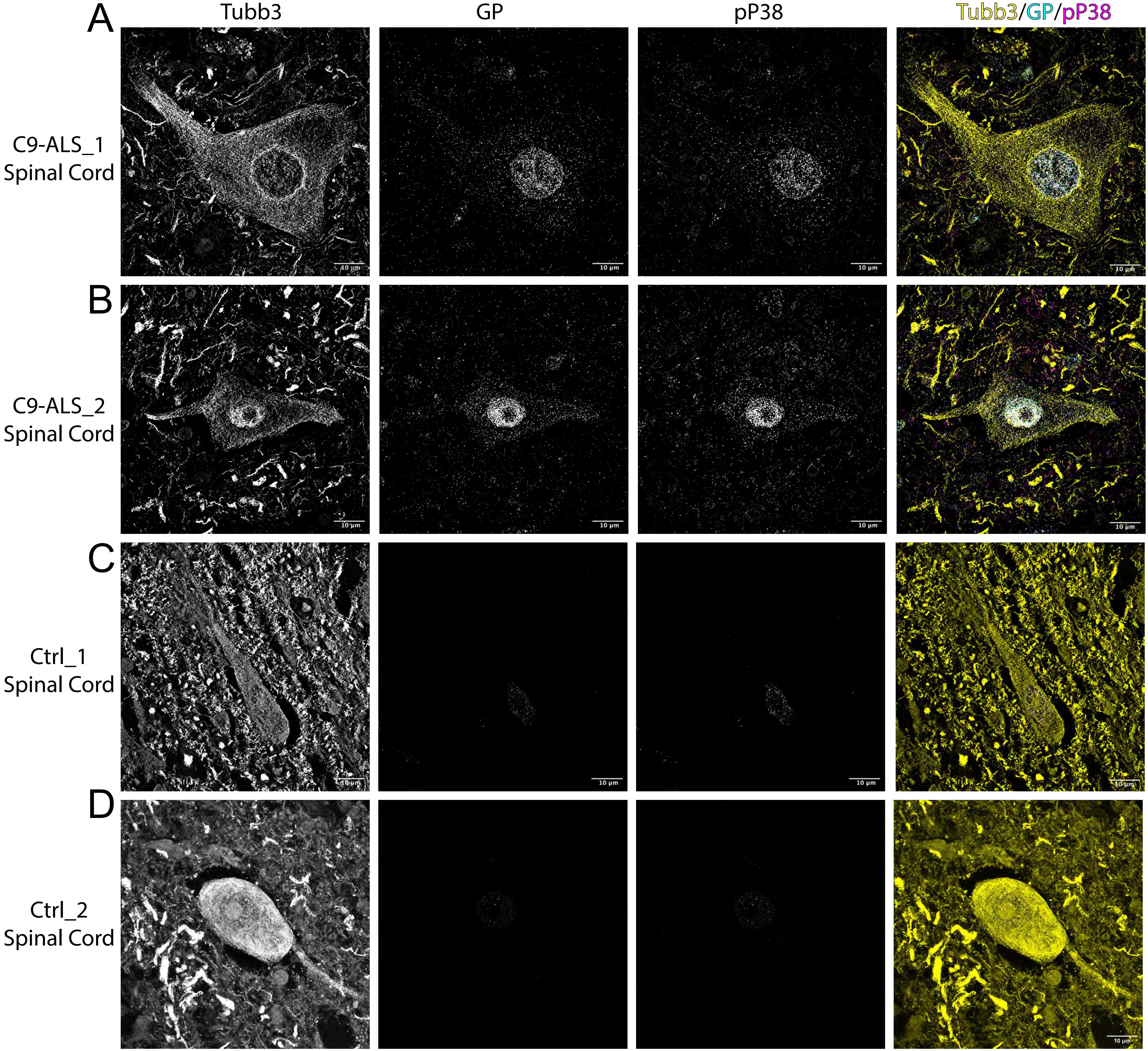
A
B
C
D
Tubb3
GP
pP38
Tubb3/GP/pP38
C9-ALS_1
Spinal Cord
C9-ALS_2
Spinal Cord
Ctrl_1
Spinal Cord
Ctrl_2
Spinal Cord
10 μm

Figure S11

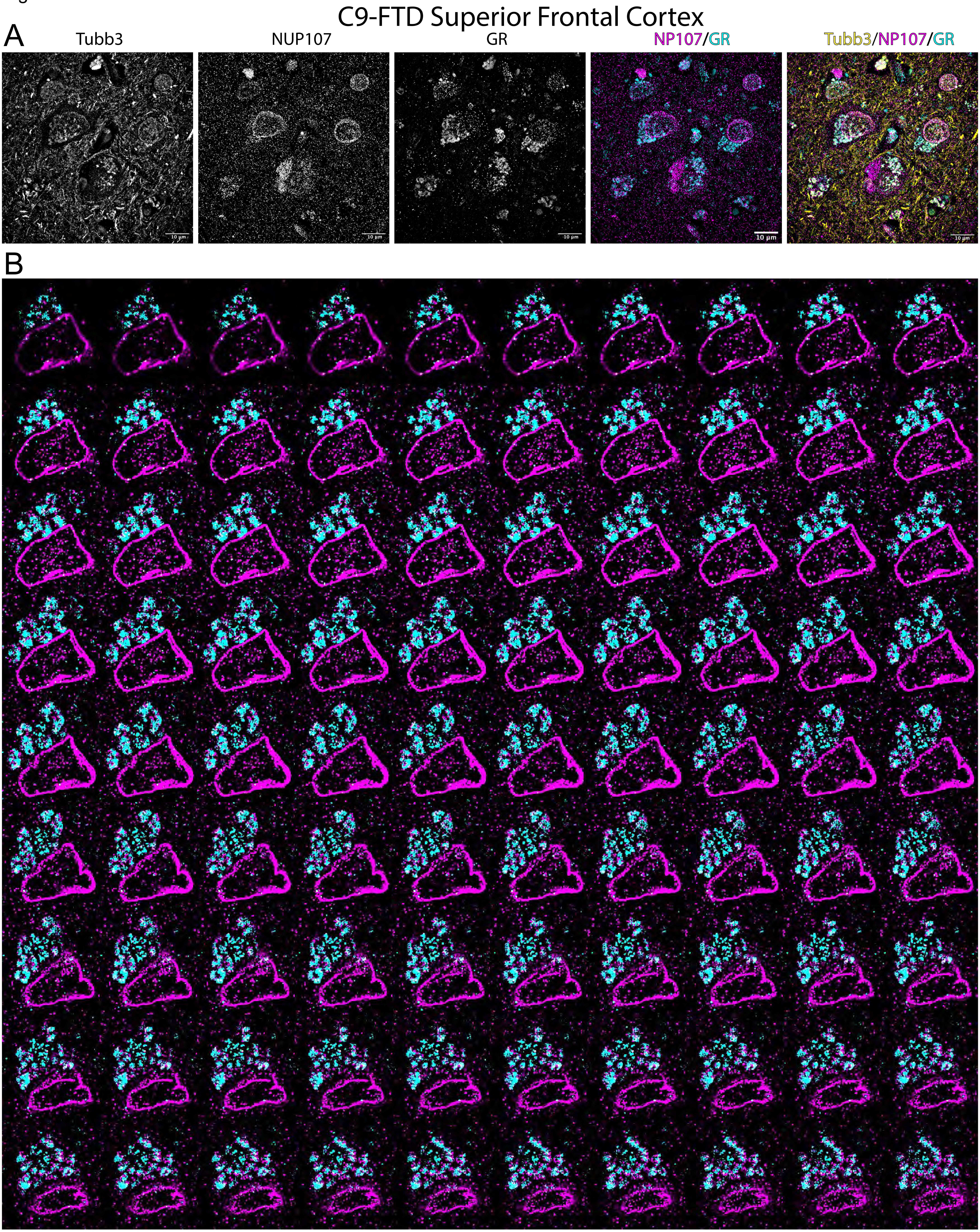
C9-FTD Superior Frontal Cortex
A
Tubb3
NUP107
GR
NP107/GR
Tubb3/NP107/GR
10 µm
B
